# CFAP61 is required for sperm flagellum formation and male fertility in human and mouse

**DOI:** 10.1101/2021.03.04.433881

**Authors:** Siyu Liu, Jintao Zhang, Zine Eddine Kherraf, Shuya Sun, Xin Zhang, Caroline Cazin, Charles Coutton, Raoudha Zouari, Shuqin Zhao, Fan Hu, Selima Fourati Ben Mustapha, Christophe Arnoult, Pierre F Ray, Mingxi Liu

**Author notes:** Corresponding authors: Mingxi Liu, Pierre F. Ray.

## Abstract

Defects in the structure or motility of cilia and flagella may lead to severe diseases such as primary ciliary dyskinesia (PCD), a multisystemic disorder with heterogeneous manifestations affecting primarily respiratory and reproductive functions. We report that CFAP61 is a conserved component of the Calmodulin and radial Spoke associated Complex (CSC) of cilia. We find that a CFAP61 splice variant, c.143+5G>A, causes exon skipping in human, inducing a multiple morphological abnormalities of the flagella (MMAF) phenotype. We generated *Cfap61* knockout mice that recapitulate the infertility phenotype of the human CFAP61 mutation, but without other symptoms usually observed in PCD. We find that CFAP61 interacts with the CSC, radial spoke stalk and RS head. During early stages of *Cfap61^−/−^* spermatid development, the assembly of RS components is impaired. With the progress of spermiogenesis, the axoneme in *Cfap61^−/−^* cells becomes unstable and scatters, and the distribution of intraflagellar transport proteins is disrupted. This study reveals an organ specific mechanism of axoneme stabilization that is related to male infertility.

## Introduction

Motile cilia and flagella are highly conserved cell organelles, they present a similar structure organized around a microtubule-based cytoskeletal called the axoneme but differ in length and functions. In mammals, defects in cilia lead to severe diseases, mainly primary ciliary dyskinesia (PCD) which affects approximately 1:10,000 individuals worldwide(Lucas et al., 2014; Rubbo & Lucas, 2017). PCD is a multisystemic disorder caused by motility defects of cilia and flagella(Afzelius & Eliasson, 1983; Munro et al., 1994), mainly characterized by recurrent respiratory tract infections with varying symptoms ranging from chronic rhinosinusitis to bronchiectasis and male infertility due to sperm immotility(Ibanez-Tallon, Heintz, & Omran, 2003). Female subfertility is less common and is caused by dysmotile fallopian tube cilia(Lyons, Saridogan, & Djahanbakhch, 2006). In 50% of cases, PCD is also associated with situs inversus, due to dysfunction of motile embryonic node cilia perturbing organ laterality(Coutton, Escoffier, Martinez, Arnoult, & Ray, 2015). More rarely, hydrocephalus arises as a consequence of ependymal cilia dysmotility leading to a blockage of the cerebrospinal fluid flow(Lyons et al., 2006). The gross axonemal structure of motile cilium and sperm tail may appear identical, but there are cell type-specific differences in axonemal proteins such as dynein arm components and in the assembly of the axoneme(Fliegauf, Benzing, & Omran, 2007). This is supported by the fact that basic motility and morphological aspects differ between motile cilia and sperm and also that not all mutations in PCD genes cause male infertility, such as CCDC114(Onoufriadis et al., 2013) and RSPH4A(Moryan, Guay, Kurtz, & Nowak, 1985). Conversely, many axonemal genes have been described to induce male infertility without any other PCD symptoms, as exemplified by *DNAH1*, which, when mutated, leads to a male infertility phenotype characterized by multiple morphological abnormalities of the flagella (MMAF)(Ben Khelifa et al., 2014). Since the first report of this severe flagellar defect in 1984, this rare phenotype has been subsequently described as ‘dysplasia of the fibrous sheath’, ‘short tails’ or ‘stump tails’ and, this heterogeneous group of flagellar defects was named ‘MMAF’ for multiple morphological anomalies of the flagella, to standardise all these terms(Coutton et al., 2015). Exome sequencing allowed researchers to partially elucidate genetic and physiopathological mechanisms that lead to sperm flagellum defects and, to date, variants in at least 19 genes have been found to be associated with the MMAF phenotype(Auguste et al., 2018; Ben Khelifa et al., 2014; Beurois et al., 2019; Coutton et al., 2019; He et al., 2019; He et al., 2020; Kherraf et al., 2018; Li et al., 2019; Li, Wu, Li, Tian, Kherraf, Zhang, Ni, Lv, Liu, Tan, Shen, Amiri-Yekta, Cazin, Liu, et al., 2020; C. Liu et al., 2019; C. Liu et al., 2020a; W. Liu et al., 2019; Wensheng Liu et al., 2019; Lorès et al., 2018; Lv et al., 2020; Martinez et al., 2020b; Martinez et al., 2018; Sha et al., 2017; Shen et al., 2019; Tang et al., 2017).

In mammals, flagella are essential for male gamete motility. The axoneme is the main component of the flagellum and runs through its entire length including the neck, middle, principle and end segments(Lindemann & Lesich, 2016). The axoneme has been extensively studied, mainly in flagella/ciliated unicellular organisms and sea urchin sperm, describing a structure conserved from single-celled protists to mammals. This core structure contains a central pair (CP) of singlet microtubules (MTs) surrounded by nine outer MT doublets. Each doublet contains a sequence of identical building blocks, which repeat longitudinally every 96 nm. Neighboring doublets are interconnected circumferentially by nexin links and are attached to the CP through radial spokes (RS). The dynein motors responsible for motility are organized into two rows of outer and inner arms along the length of the doublets. Dyneins are minus end–directed motors, and their unidirectional movement along a B-tubule induces bending in one direction, whereas the dyneins located on the opposite side of the axoneme induce bending to the opposite direction(Heuser, Dymek, Lin, Smith, & Nicastro, 2012). Structures important for the coordination of dynein activity include the CP complex, the RSs, the I1 inner arm dynein, and the dynein regulatory complex (DRC)(Coutton et al., 2018; Dong et al., 2018; Kherraf et al., 2018; Lorès et al., 2018; Sha et al., 2017).

Björn Afzelius, in 1959, was the first to describe the presence of RSs in the axonemes of sea urchin sperm flagella(Satir, 1968). In addition to their structural role in maintaining the 9+2 axoneme stability, RSs seem to also be involved in the beating motion, participating in signal transduction between the CP and the dyneins, (F. Abbasi et al., 2018; C. Liu et al., 2020b; Shinohara et al., 2015). Initially, SDS-PAGE analysis of axonemes of wild-type (WT) and paralyzed mutants of Chlamydomonas revealed 17 polypeptide chains that were ascribed to the RS complex(Piperno, Huang, Ramanis, & Luck, 1981; Whitfield et al., 2019), and the eventual purification of the RS complex (Beurois et al., 2019; Li, Wu, Li, Tian, Kherraf, Zhang, Ni, Lv, Liu, Tan, Shen, Amiri-Yekta, Cazin, Liu, et al., 2020) enabled the identification of 23 proteins(Beurois et al., 2019; Martinez et al., 2020a). These proteins were called RSP1 to RSP23 and, in Chlamydomonas, the RSPs 1–12, 14, 16-17, 20, 22 and 23 have been identified and sequenced(Curry, Williams, & Rosenbaum, 1992; King & Patel-King, 1995; Patel-King, Gorbatyuk, Takebe, & King, 2004; Williams, Velleca, Curry, & Rosenbaum, 1989; C. Yang, Compton, & Yang, 2005; P. Yang et al., 2006; P. Yang, Yang, & Sale, 2004; Zimmer, Schloss, Silflow, Youngblom, & Watterson, 1988). RSPs are assembled into RSs in two phases. Separation of the RSPs on sucrose density gradients permitted to identify and characterize different S(ucroses) fractions/particles. First, the cell assembles partial RSs as 12S particles, composed of RSPs 1-7 and 9-12(Diener et al., 2011; Pigino et al., 2011; Qin, Diener, Geimer, Cole, & Rosenbaum, 2004). After delivery to the flagella by intraflagellar transport (IFT), 12S precursors are converted into 20S mature RSs, by the assembly of the remaining RSPs(Diener et al., 2011; Pigino et al., 2011) Regarding these later assembled RSPs, the knowledge on RSPs 13, 15, 18, 19, 21 is still relatively limited, but interestingly, in subsequent studies, RSP18 and RSP19 were found to be involved in the calmodulin (CaM)- and spoke-associated complex (CSC). Both RSP18 and RSP19 are present in axonemes from pf14, a Chlamydomonas radial spoke mutant that lacks radial spoke structure(Piperno et al., 1981).

In the present study, we found that a splice-site variant c.143+5G>A of the Rsp19 homologous gene, CFAP61, leads to exon skipping and induces MMAF in humans. In addition, CFAP61 has been found to interact with the CSC, RS stalk and RS head. Lack of *Cfap61* in mice resulted in an MMAF phenotype and male sterility. In the early stages of *Cfap61^−/−^* spermatid development, RS 12S precursors were assembled, but the assembly of other RS components was blocked and, with the progress of spermiogenesis, the axoneme became unstable and was severely altered in *Cfap61* knockout mice. This defect was only observed in the assembly of the flagellum axoneme, while there was no effect on cilia. In addition, the absence of *Cfap61* also affected the distribution of IFT in the sperm flagellum. Therefore, this study reveals an organ dependent mechanism of axoneme stabilization that is related to male infertility.

## Results

### *Cfap61* is an evolutionarily conserved gene predominantly expressed in the testis

In Chlamydomonas, the CSC is located at the base of RS and interacts with the DRC and inner dynein arm (IDA)(Dymek & Smith, 2007; Heuser et al., 2012; Urbanska et al., 2015; Viswanadha, Sale, & Porter, 2017) (Figure 1A), while CaM interacting proteins, CaM-IP1-3 (FAP91, FAP61 and FAP251), are considered components of the CSC(Heuser et al., 2012). Mutations in the human homologues, *MAATS1* (FAP91 homologous gene)(Martinez et al., 2020a) and *CFAP251* (FAP251 homologous gene)(Auguste et al., 2018; Kherraf et al., 2018; Li et al., 2019) have been detected in patients with MMAF. We evaluated the expression of potential CSC components in various organs in mice and found that CSC components were predominantly expressed in the testis (Figure 1B-D). The FAP61 homologous gene, *Cfap61*, was detectable in testis, brain and lung, but its testis expression level was much higher than that in other organs. *Maats1* (FAP91 homologous gene) and *Wdr66* (FAP251 homologous gene) showed similar expression patterns in mice. Regarding the components of RS, we observed a distinct expression pattern in different organs. For example, *Armc4* and *Rsph9* are preferentially expressed in the testis, while *Rsph4a* is highly expressed in the lung (Figure 1E-G). Among basal eukaryotes, *Cfap61* is expressed in species that present a flagellum at some stage of the organism’s life cycle. The CFAP61 protein is composed of a domain of unknown function (DUF4821) and protein sequence alignment showed that CFAP61 is an evolutionarily conserved gene present among human, mouse, rat, Xenopus, zebrafish and Chlamydomonas (Figure 1H).

**Figure 1:**
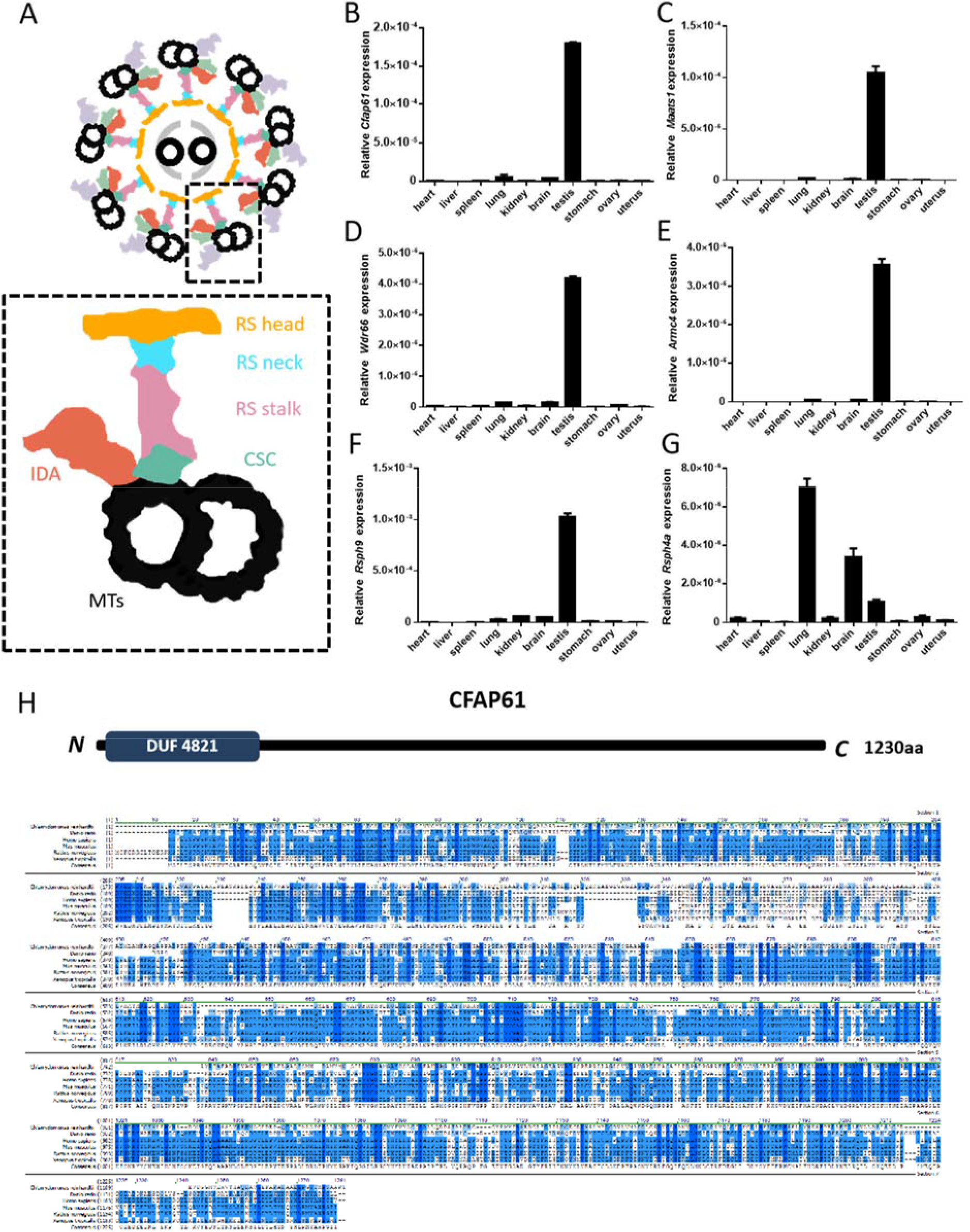
*Cfap61* is an evolutionarily conserved testis-enriched gene. (A) Schematic drawing of the flagella axoneme cross-section. Microtubules (MTs), inner dynein arm (IDA), radial spoke (RS) components and Calmodulin (CaM) and spoke associated complex (CSC) are indicated in the dotted box. (B-G) Quantitative RT–PCR results showing relative expression levels of *Rsph* and CSC genes in several mouse organs. (H) Sequence similarity of CFAP61 protein in various organisms. Dark blue background indicates identical residues in all species while blue background represents conserved residues. Light blue background shows weakly similar residues.

### Exome sequencing identified CFAP61 homozygous variants in patients with MMAF

A cohort of 167 patients with MMAF and no other signs of PCD was previously analyzed and permitted to identify harmful variants in known MMAF-related genes in 66 patients (Martinez et al., 2020a). After reanalysis of the exomes data from the undiagnosed subjects, we identified one patient with a homozygous variant in CFAP61. This patient (PaCFAP61) had an intronic variant predicted to alter splicing: c.143+5G>A (NM_015585.4) (Figure 2A). The variant was present in the Genome Aggregation Database with a minor allele frequency of 1.59e-5. Additionally, we found by minigene assay that c.143+5G>A induced the skipping of CFAP61exon 2 (151nucleotides) and induced a frame-shift shortly after exon1 (Figure 2B and C). These results suggest that c.143+5G>A affects the normal function of CFAP61 in humans and leads to MMAF. PaCFAP61 presented a typical MMAF phenotype with a semen volume of 3ml, 40M of sperm/ml, 20% of progressive motility and multiple flagellar anomalies: 36% of bent, 18% with no tail, 12% with a short tail, 50% with an irregular shape and 30% with a coiled tail.

**Figure 2:**
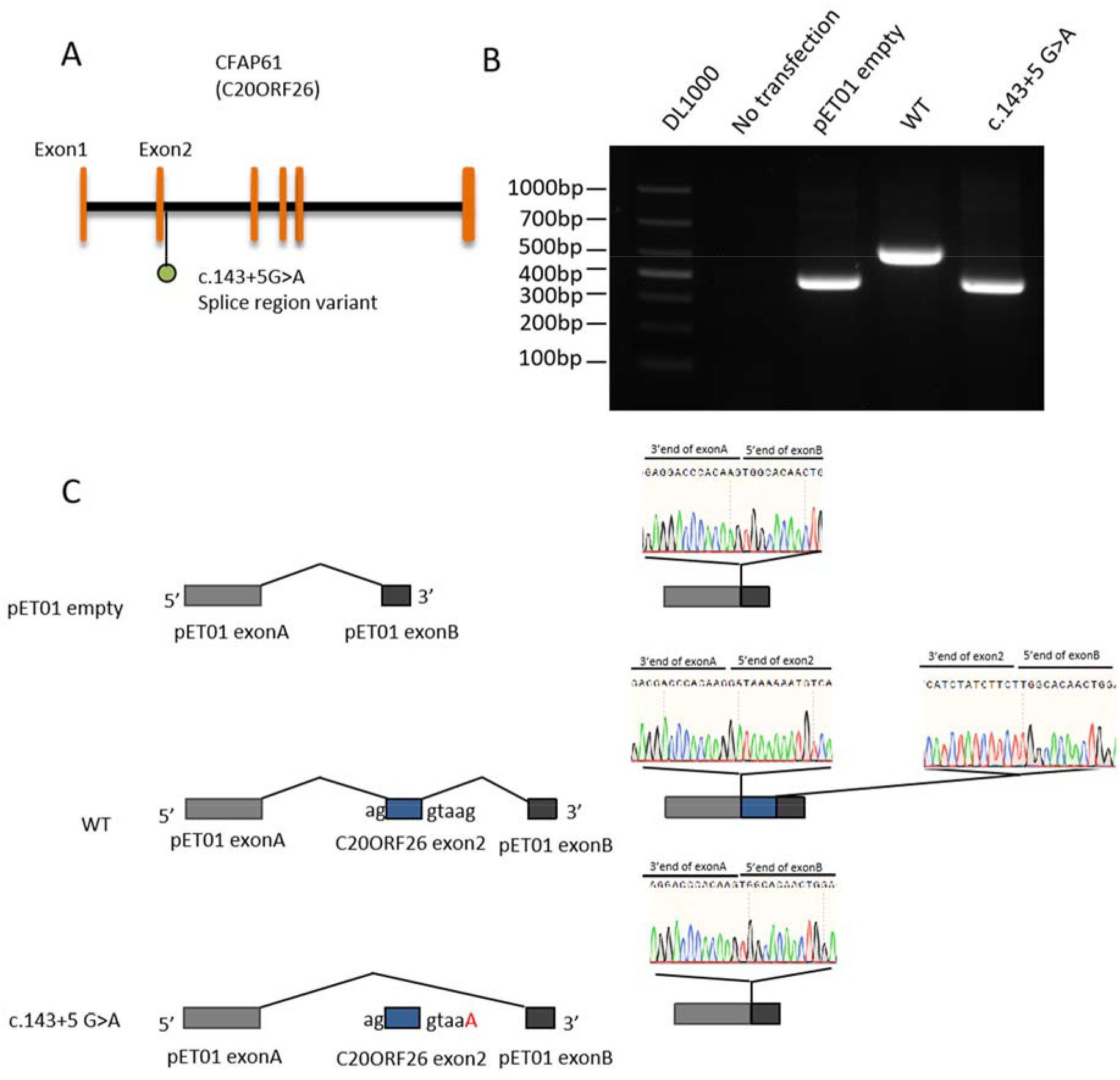
Results of the minigene assay and splicing impact of the c.143+5 G>A mutation on *Cfap61*. (A) A homozygous splice region variant: ENST00000245957.10:c.143+5G>A. Electrophoregrams of Sanger sequencing for PaCFAP61 and Control. (B) Gel electrophoresis that c.143+5 G>A causes abnormal mRNA splicing. (C) Sanger sequencing confirmed that the c.143+5 G>A variant causes complete exon2 skipping.

### CFAP61 is required for male fertility and sperm flagellum formation

To examine the function of *Cfap61* in vivo, we generated a *Cfap61* mutant mouse strain using the CRISPR/Cas9 system. A stable *Cfap61* mutant mouse line carrying two allelic variants: a 1-bp and 10-bp deletion within exon4 (Figure 3A); was established and male mice homozygous for this allele are referred herein as *Cfap61^−/−^*. Two mouse CFAP61 peptides were selected as antigens to generate specific antibodies (Figure 3-figure supplement 1) and western blot and immunofluorescence confirmed that CFAP61 was completely erased in *Cfap61^−/−^* testis (Figure 3B and C). In testicular histological sections, CFAP61 was mainly distributed in the spermatozoa in seminiferous tubules from wild-type mice (Figure 3C). *Cfap61^−/−^* males show no overt abnormality in development or behavior. To test male fertility, individual males (*Cfap61^+/+^* and *Cfap61^−/−^*) were housed with *Cfap61^+/+^* (wild-type) females and the number of pups per litter was recorded. *Cfap61^−/−^* males failed to sire any offspring despite copulating with females (Figure 3D), demonstrating that *Cfap61* is required for male fertility.

**Figure 3:**
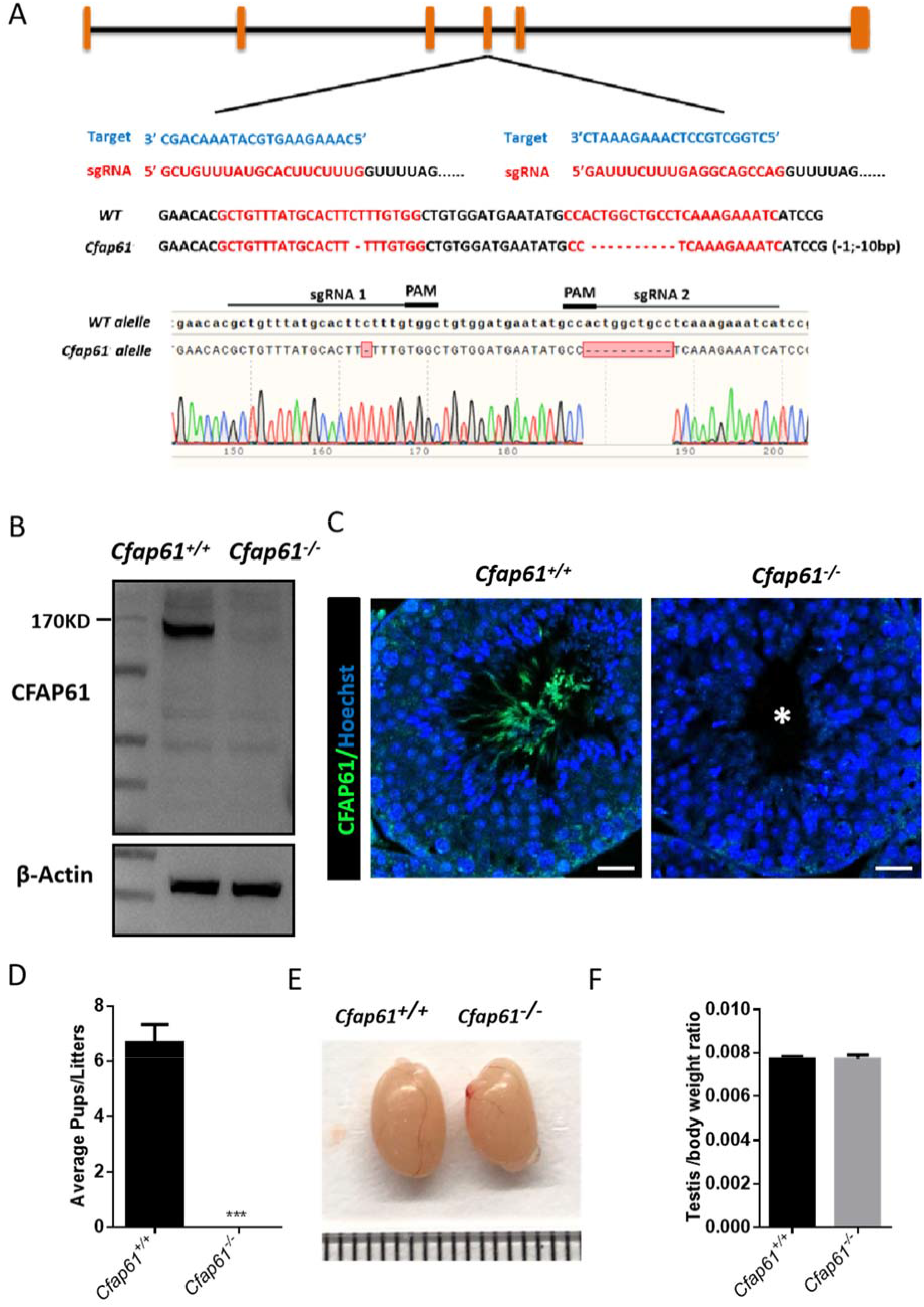
Cfap61−/− mice are infertile. Schematic diagram of CRISPR/Cas9 targeting strategy. The sgRNAs were designed within exon 4 of *Cfap61* and a 1-bp and 10-bp deletion was detected in *Cfap61^−/−^* mice by Sanger sequencing. (B, C) CFAP61 was not detected in *Cfap61^−/−^* testis by western blot (B) and immunofluorescence (C). Scale bars =20 μm. (D) The average litter size of WT (6.677± 0.667), *Cfap61^−/−^* (0) male mice (n=3).P (*Cfap61^−/−^* vs WT)= 0.0006. (E) The size of the testes was not altered in the WT and *Cfap61^−/−^* adult mice. (F) The testis/body weight ratio of WT (0.007689±0.0001418), *Cfap61^−/−^* (0.007721±0.000175) male mice (n=3).P (*Cfap61^−/−^* vs WT)= 0.8948. Data represent mean±SEM.

To determine if a block in spermatogenesis was the reason for infertility, we examined spermatogenesis in *Cfap61^−/−^* males. Gross examination of testis revealed no difference in appearance and testis weight between *Cfap61^+/+^* and *Cfap61^−/−^* littermates (Figure 3E and F). Using Periodic acid–Schiff (PAS) staining, we found that *Cfap61^−/−^* lacked flagella with normal length in the lumen of seminiferous tubules (Figure 3-figure supplement 2). Western blot analysis showed that CFAP61 was mainly expressed in the testis among ciliated/flagellated organs (Figure 4A) and protein extracts from sperm revealed that CFAP61 is in a Triton-resistant, SDS-soluble pool (Figure 4B). Confocal microscopy reveals CFAP61 expression along the flagella in mouse spermatids (Figure 4C-N).

**Figure 4.**
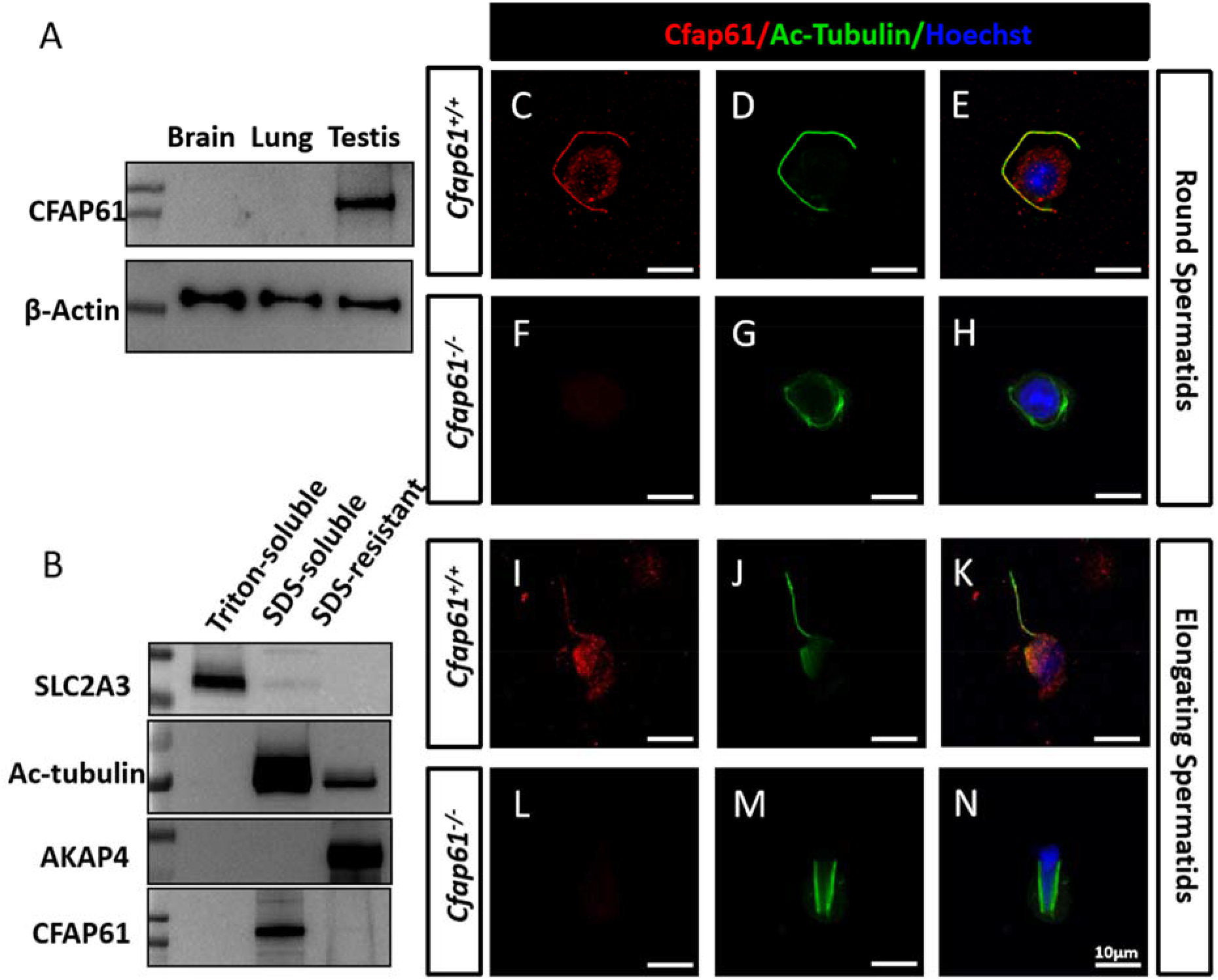
CFAP61 localizes to sperm tails. (A) Western blot analysis of protein extracts from brain, lung, and testis. (B) Western blot analysis of sperm fractionated into Triton X-100 soluble, SDS soluble, and SDS insoluble fractions from wild-type mice. SLC2A3, acetylated tubulin and AKAP4 were detected as markers for Triton-soluble, SDS-soluble and SDS-resistant fractions, respectively. (C-N) Immunofluorescence analysis of round and elongating spermatids from wild-type and *Cfap61^−/−^* mice using anti-CFAP61 (red) and anti-acetylated tubulin (green) antibodies. Scale bars =10 μm.

When examining epididymis sections, *Cfap61^−/−^* seemed to contain less spermatozoa in the cauda and caput regions than *Cfap61^+/+^* mice (Figure 3-figure supplement 2), which was confirmed by the number of sperm collected from epididymal cauda of knockout mice that was significantly lower than in wild-type mice (Figure 5A). Almost all *Cfap61^−/−^* spermatozoa were abnormal and showed short, bent, curled, thick or missing flagella (Figure 5-figure supplement 1). Similar phenotypes were reported in a contemporaneous study of another *Cfap61* knockout mouse(Huang et al., 2020). The acrosome morphology of *Cfap61^−/−^* sperm was normal, while the formation of the tail was clearly disordered (Figure 5B-F). These severe tail deformities of *Cfap61^−/−^* sperm have great influence on sperm movement. *Cfap61^−/−^* sperm showed a state of immobility (Movie 1 and Movie 2). Taken together, these observations suggest that *Cfap61* is required for male fertility and sperm flagellum formation and beating.

**Figure 5.**
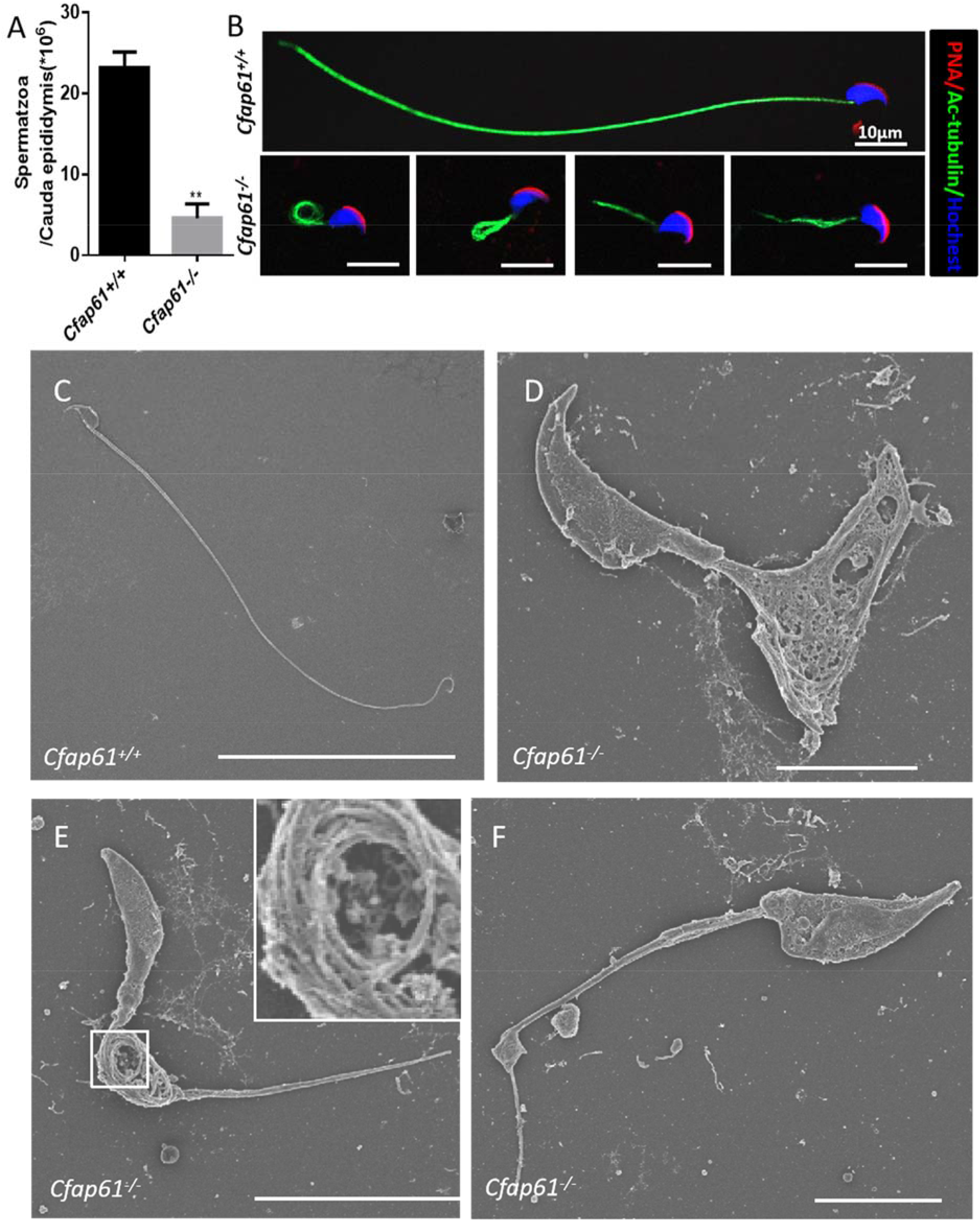
Spermatozoa appear abnormal in *Cfap61^−/−^* mice. (A) The sperm concentration of WT (23.15±1.998) and *Cfap61^−/−^* (4.593±1.760) male mice (n=3).P (*Cfap61^−/−^* vs WT)= 0.002. Data represent mean±SEM. (B) Fluorescence detection of AC-tubulin (green) and PNA (red) on wild-type and *Cfap61^−/−^* spermatozoa. (B)Scale bars =10 μm.(C-F) Scanning electron micrographs of wild-type (C) and *Cfap61*-knockout (D-F) sperm. The arrangement of microtubules was reticular (D), coil like (E) and irregular in different segments (F). The enlarged section shows coil like arrangement of microtubules (E). Sperm from wild-type mice show normal morphology while sperm from *Cfap61^−/−^* mice show severe flagella morphology defects. (C)Scale bar =50 μm.(D, F) Scale bars =5 μm.(E) Scale bar =10 μm.

### CFAP61 is a component of sperm flagella CSC and radial spoke

We next performed immunoprecipitation (IP)-mass spectrometry of CFAP61 to determine the CFAP61 interactome in mouse testis (Figure 6-figure supplement 1A). We found that the proteins interacting with CFAP61 include axonemal components and involved in its functional regulation and assembly (Figure 6-figure supplement 1B). Through co-IP, we confirmed that murine CFAP61 interacts with MAATS1 from CSC (Figure 6A and B). CFAP61 can also interact with RS stalk proteins, such as ARMC4 (RSPH8), RSPH3A and ROPN1/ROPN1L (RSPH11) (Figure 6C-F). A RSP22 homologous protein, DYNLRB2, was detected in interaction with CFAP61 by mass spectrometry (Figure 6-figure supplement 1B), but co-IP results showed that CFAP61 and DYNlRB2 (RSPH22) had no direct interaction with each other (Figure 6-figure supplement 2A). These results suggest that any interaction between CFAP61 and DYNlRB2 in the testis would be indirect.

**Figure 6.**
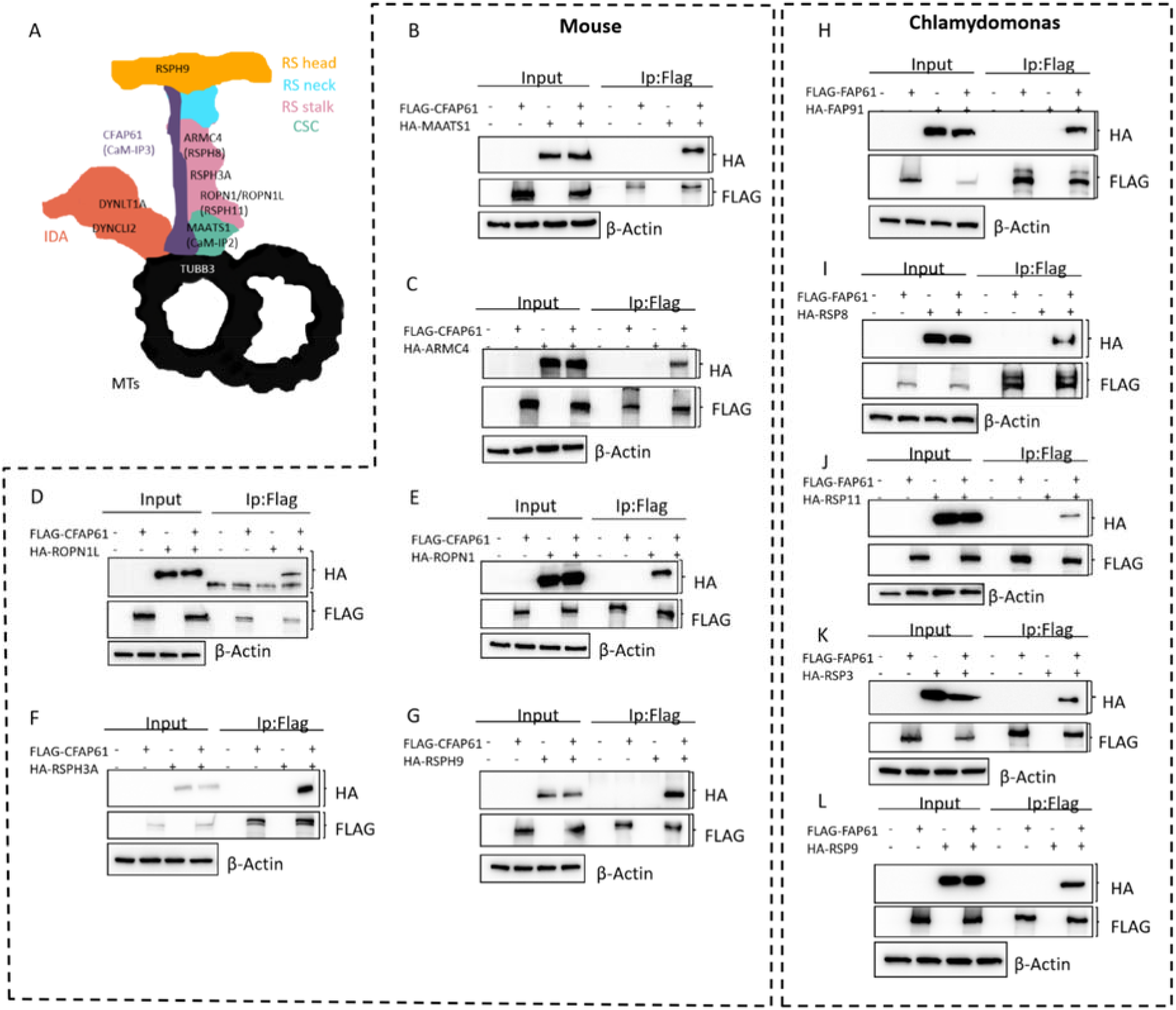
CFAP61 is a component of radial spoke and interacts with other proteins. (A) Diagram of CFAP61 localization within the radial spoke and CSC. (B-L) Mouse and Chlamydomonas RS or CSC components were expressed or co-expressed with CFAP61 in HEK293T cells and CFAP61 interaction with other RS and CSC proteins was examined by co-immunoprecipitation with mouse (B-G) or Chlamydomonas (H-L) proteins.

Since previous reports have suggested that CFAP61 is a CSC component(Dymek & Smith, 2007; Heuser et al., 2012; Urbanska et al., 2015), an interaction between CFAP61 and the RS stack can be predicted. Surprisingly, both our mass spectrometry data and co-IP data showed that CFAP61 can also interact with RSPH9, the RS head component (Figure 6-figure supplement 1B and Figure 6G). Similar to data in mice, the co-IP assay showed that Chlamydomonas fap61 also interacted directly with FAP91, RSP8, RSP3, RSP11 and RSP9 (Figure 6H-L), but not with RSP22 (Figure 6-figure supplement 2B). In addition, we found that CFAP61could interact with DYNCLI2 and DYNLT1A of the dynein arm and TUBB3 (Figure 6-figure supplement 2C-E). In Chlamydomonas, the dynein arm IA4 (dynein E) is greatly reduced in all CSC mutants(Heuser et al., 2012). These results suggest that CSC interaction with the dynein arms is conserved among species.

### Deletion of CFAP61 resulted in abnormal sperm flagella RS, but did not affect the structure and function of respiratory cilia

In order to investigate whether CFAP61 deletion affects the assembly of RS, we used confocal microscopy to access the expression of RSPH9 and NME5 (RSPH23) proteins in sperm. Both RSPH9 and NME5 signals were very low in *Cfap61^−/−^* spermatozoa (Figure 7A-J). To investigate if *Cfap61^−/−^* mice recapitulated the respiratory phenotype observed in PCD, we evaluated the distribution of RS proteins in tracheal cilia. Immunofluorescence signals of RSPH9 and NME5 in trachea cilia were evaluated by high resolution microscopy, and no difference was observed between *Cfap61^−/−^* and *Cfap61^+/+^* mice (Figure 8A and B). Additionally, no expression of CFAP61 was detected in the respiratory tract cilia in both *Cfap61^−/−^* and *Cfap61^+/+^*mice, which is consistant with the western blot results (Figure 6A). Scanning electron microscopy did not detect any significant change in the length of respiratory tract cilia in *Cfap61^−/−^* mice (Figure 8C-F). Similarly, there was no difference in the beating of trachea cilia from *Cfap61^−/−^* and *Cfap61^+/+^* animals (Movie 3 and Movie 4). These results suggest that CFAP61 plays an essential role in the formation of the RS structure in mammalian sperm flagellum, but is not necessary for respiratory cilia.

**Figure 7.**
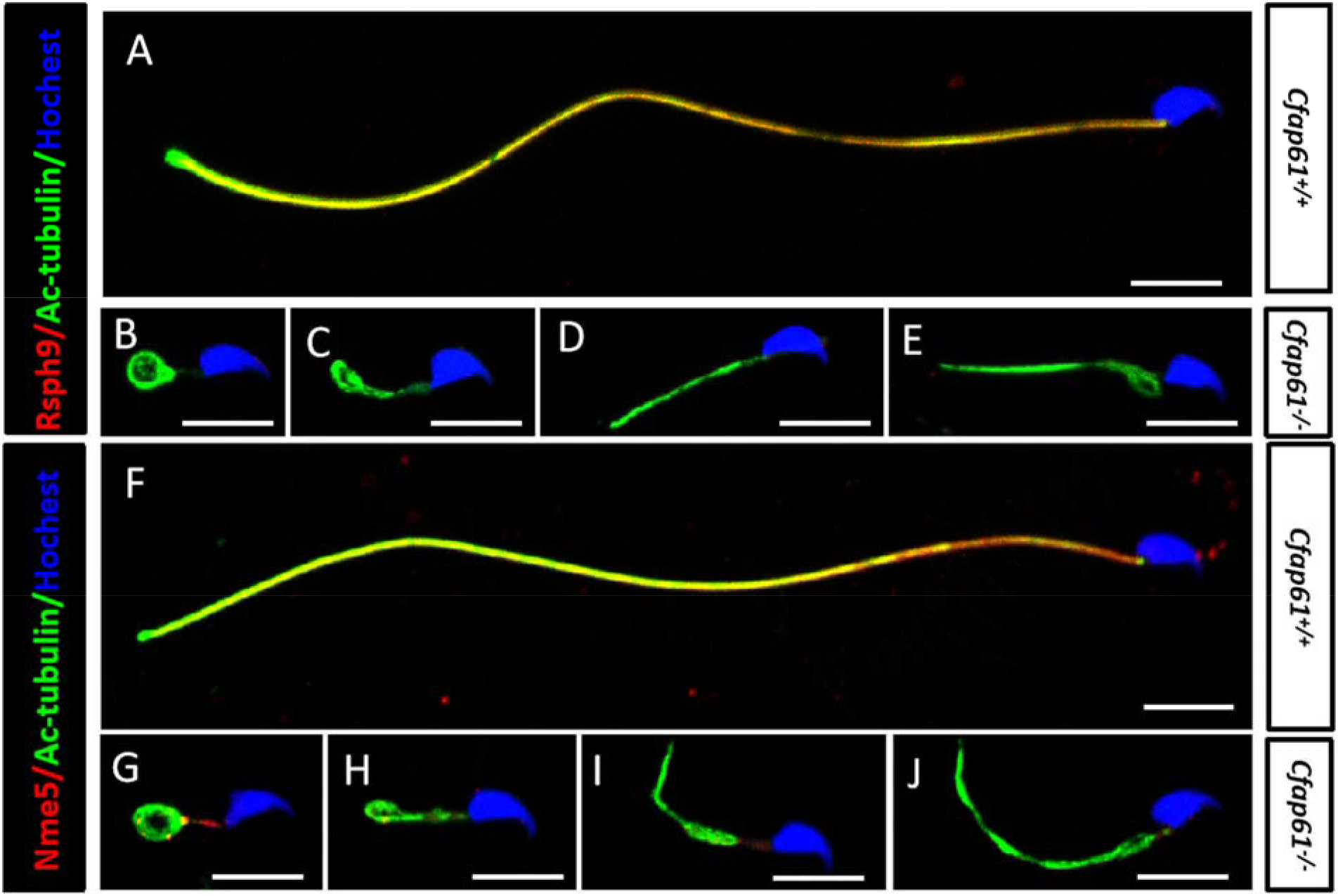
Immunofluorescence analysis of radial spoke proteins in mouse sperm flagellum. (A-E) Subcellular localization of RSHP9 (ref) and acetylated tubulin (green) in the sperm flagellum in wild-type (A) and *Cfap61^−/−^* mice (B-E). (F-J) Subcellular localization of NME5 (red) and acetylated tubulin (green) in the sperm flagellum in wild-type (F) and *Cfap61^−/−^* mice (G-J).Scale bars =10 μm.

**Figure 8.**
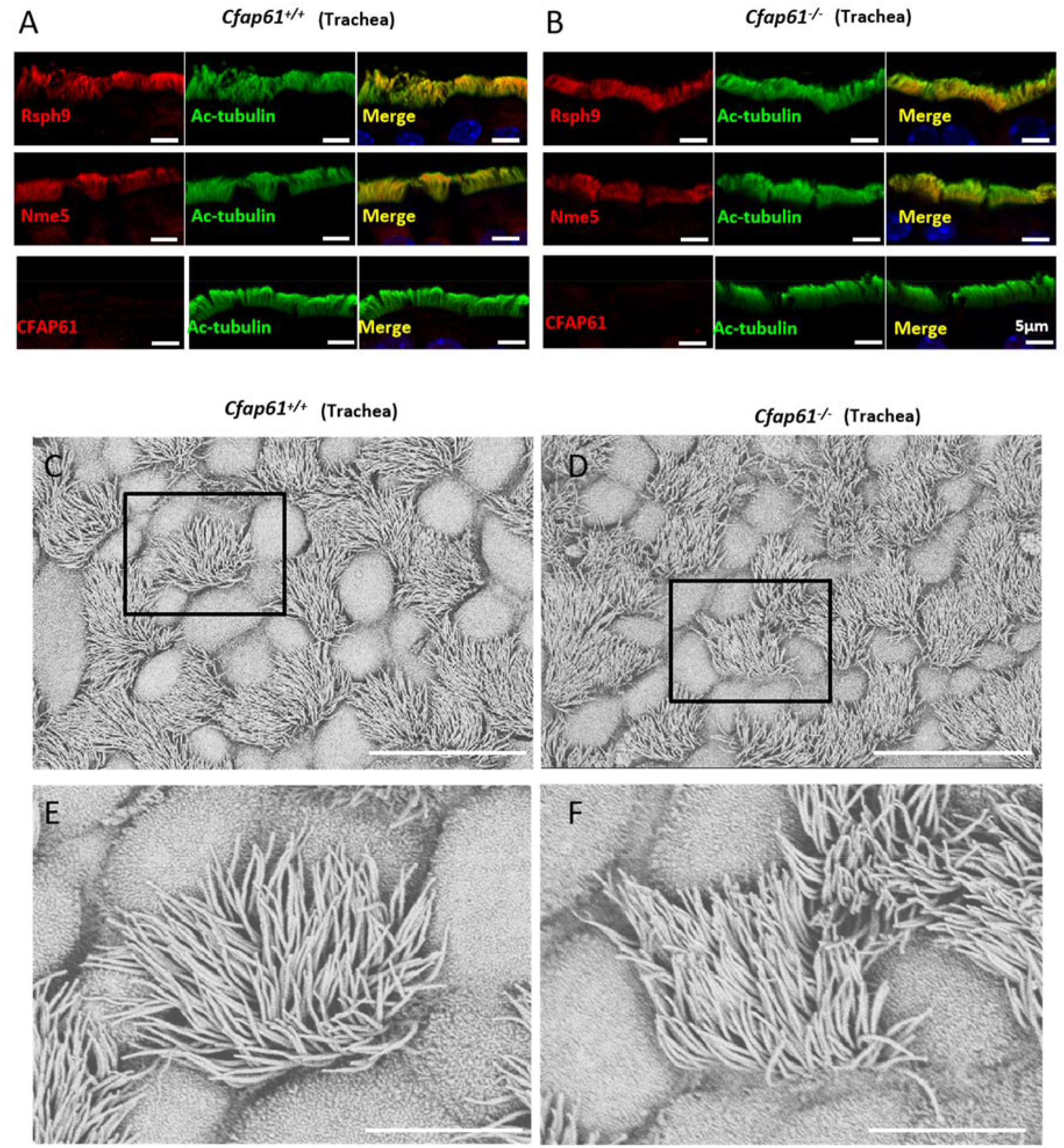
Trachea cilia appear normal in *Cfap61^−/−^* mice. (A,B) Subcellular localization of RSHP9, NME5, CFAP61 (red) and acetylated tubulin (green) in trachea cells of wild-type (A) and *Cfap61^−/−^* mice (B). (C-F) Scanning electron micrography of wild-type (C, E) and *Cfap61^−/−^* (D, F) tracheal epithelium at low (C, D) and high magnifications (E, F). (A, B) Scale bars =5 μm. (C, D) Scale bars =20 μm. (E, F) Scale bars =60 μm.

### The absence of *Cfap61* leads to flagella axoneme assembly failure and sperm deformation

In order to understand the role of CFAP61 for flagellum assembly, we analyzed flagellum formation during spermiogenesis (Figure 9A-O) and found that in *Cfap61^−/−^* mice, axonemes formed normally until the step 2 of spermatid formation (9G). However, with the progress of spermatid differentiation, the microtubule structure became disordered and could not form a normal axonemal structure (Figure 9H-O).Using transmission electron microscopy, we observed that *Cfap61^−/−^* microtubules and outer dense fibers were separated from the axoneme (Figure 9P and Q); however, centriole anchoring and implantation fossa formation were not affected (Figure 9R). We further assessed the assembly of RS components during spermiogenesis. In the axoneme of Chlamydomonas, the 12S RS complex as a whole is assembled first, and the other RS components, such as RSP23 and RSP16, are assembled later as the 20S RS complex (Diener et al., 2011; Pigino et al., 2011). In mice, we found that the assembly of RSPH9 in round spermatids was normal (Figure 10A-D), but was missing in elongating spermatids of *Cfap61* knockout mice (Figure 10E-J). In addition, we show that the non-12S RS component, NME5 (RSPH23), failed to assemble in *Cfap61^−/−^* round and elongating spermatids (Figure 10K-T). With the genesis of sperm morphological deformation, the axoneme become unstable and completely scattered. This suggests that CFAP61 plays an irreplaceable role in the assembly of the 20S RS complex in mammalian flagella and indicates that incomplete assembly of 20S RS component does not allow RS stability. In the late stages of spermatid flagellum assembly, the assembled 12S RS proteins seem to disappear from the flagellum due to the absence of CFAP61. Interestingly, there was no RS assembly abnormality in *Cfap61^−/−^* respiratory cilia, suggesting that *Cfap61* function is limited to sperm flagellum in mammals.

**Figure 9.**
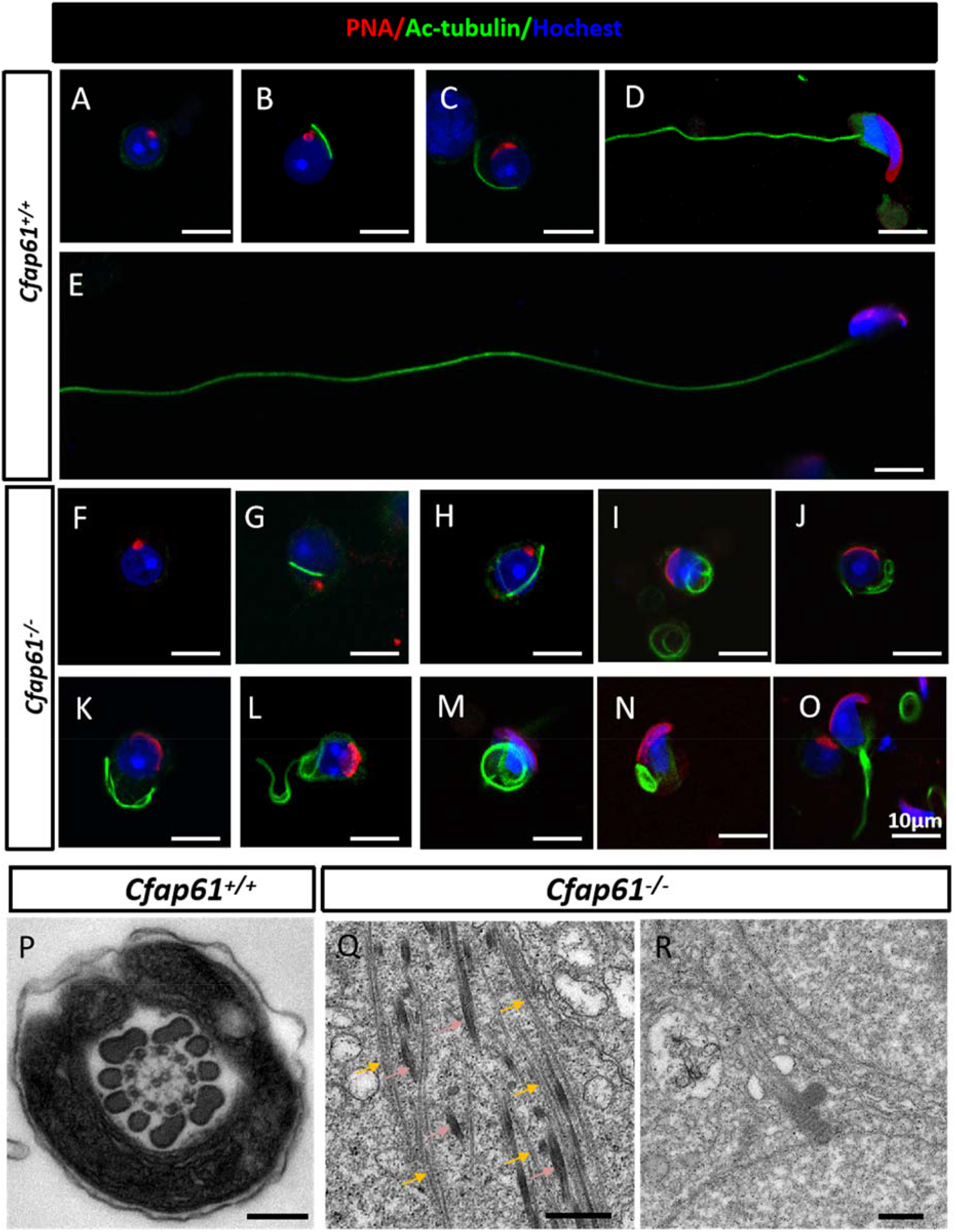
Abnormal sperm flagella assembly of *Cfap61^−/−^* mice occurs from step 3 round spermatids. (A-O) Immunofluorescence analysis of acetylated-tubulin (green) and PNA (red) from wild-type (A-E) and *Cfap61^−/−^*germ cells (F-O). Ultrastructural analysis of wild-type (P) and *Cfap61^−/−^* testicular spermatozoa flagella (Q, R). (Q)The dispersed microtubules could not form “9 + 2” axonemes. The yellow arrow indicates microtubule. The pink arrow indicates the outer dense fiber. (R)The centriole of round sperm can be anchored to the nuclear membrane. (A-O) Scale bars =10 μm.(P, Q) Scale bars =200 nm.(R) Scale bar =500 nm.

**Figure 10.**
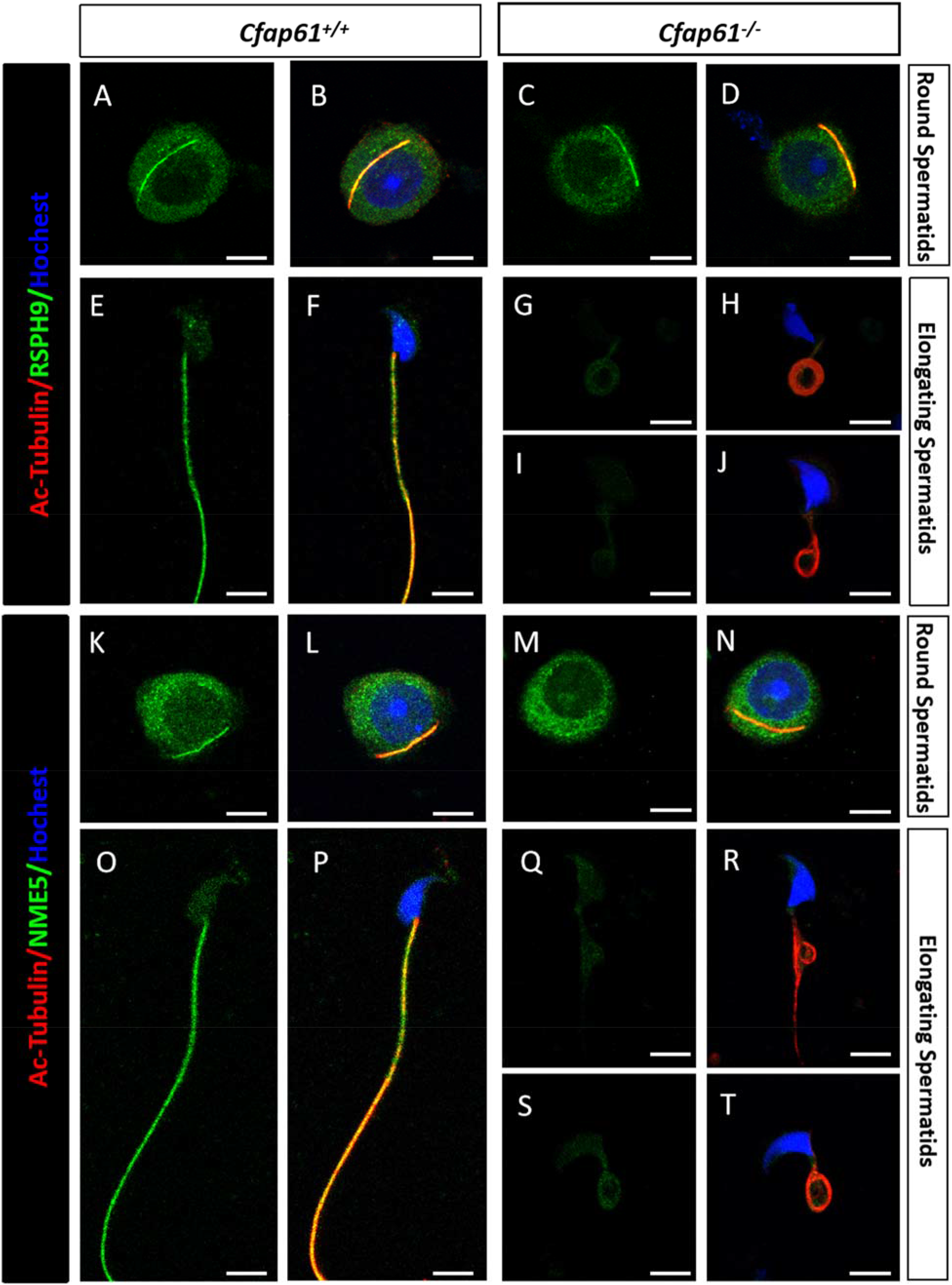
Ablation of *Cfap61* affects the assembly of 20S radial spoke complex, while the 12S complex is unaffected. (A-J) Immunofluorescence analysis of acetylated-tubulin (red)and RSPH9 (a component of the 12S radial spoke complex; green) in wild-type (A, B, E, F) and *Cfap61^−/−^* germ cells (C, D, G, H, I, J). (K-T) Immunofluorescence analysis of acetylated-tubulin (red) and NME5 (a component of the 20S radial spoke complex; green) in wild-type (K, L, O, P) and *Cfap61^−/−^* germ cells (M, N, Q, R, S, T). Scale bars =5 μm.

Through co-immunoprecipitation, we demonstrate that CFAP61 can interact with intraflagellar transport (IFT) components, such as WDR35, IFT22 and IFT81 (Figure 11A-C). Although we did not observe direct interaction between CFAP61 and IFT74 (Figure 11D), IP-mass spectrometry showed that CFAP61 and IFT74 may interact indirectly, as IFT74 is a component of the IFT-B subcomplex(Lechtreck, 2015). Immunofluorescence showed that the IFT74 structure was located in the axoneme at the early stage of flagellum assembly, but not in the axoneme of elongating spermatid, when it was mainly located in the manchette (Figure 11E-G). Similar to IFT74, CFAP61 is also located in the manchette of the elongating spermatid (Figure 4I-K). In *Cfap61^−/−^* spermatids, no changes were observed in IFT proteins in the early stage of spermiogenesis, but they appeared to be trapped in the flagellum of the elongating spermatid (Figure 11H-L). Although we are not sure whether this IFT retention is another cause for the abnormal flagellum assembly, it shows that the absence of *Cfap61* has a variety of effects on flagellum formation.

**Figure 11:**
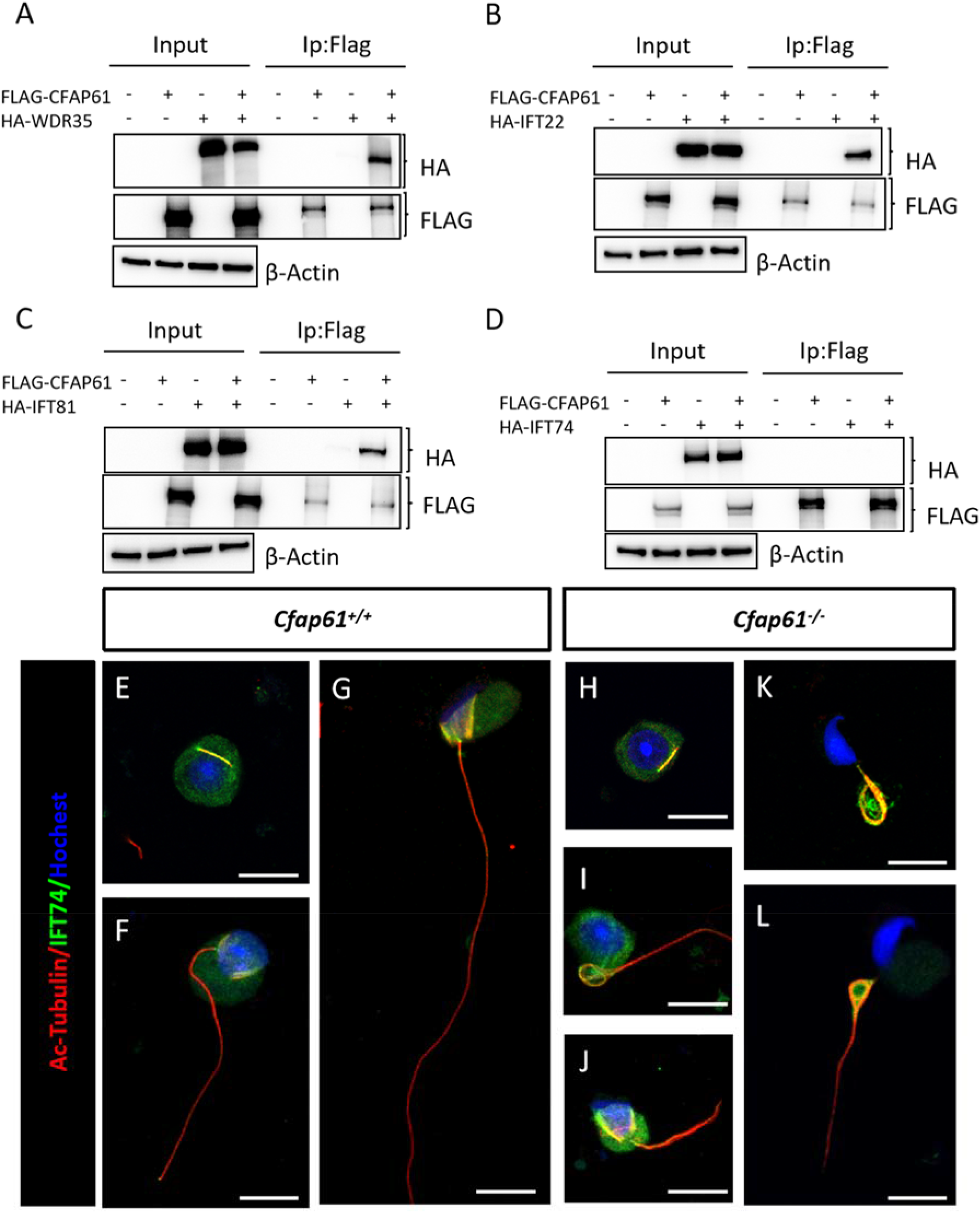
Intraflagellar transport protein retenet in mature spermatozoa. (A-D) Intraflagellar transport (IFT) proteins were expressed or co-expressed with CFAP61 in HEK293T cells, and CFAP61 interaction with IFT proteins was examined by co-immunoprecipitation. (E-L) Immunofluorescence analysis of acetylated-tubulin (red) and IFT74 (green) in wild-type (E-G) and *Cfap61^−/−^* germ cells (H-L). Scale bars =10 μm.

## Discussion

In the present study, we describe a novel CFAP61 variant c.143+5G>A inducing exon skipping and participating in the pathogenesis of MMAF. In Chlamydomonas, the CFAP61 homologous protein FAP61 was initially detected in the 20S RS complex(Beurois et al., 2019; Martinez et al., 2020a), but in later studies FAP61 was found to remain in RS1 and RS2 deleted flagella. Meanwhile, FAP61 can interact with CaM, suggesting that it may be a CSC component(Dymek & Smith, 2007; Heuser et al., 2012; Urbanska et al., 2015). Our study found that CFAP61 is a CSC component and also a RS component, and that CFAP61 can interact with CSC components and with ARMC4 (RSPH8), RSPH3, AROPN1/ROPN1L (RSPH11) and RSPH9 of the RS complex. These interactions also exist in Chlamydomonas homologous proteins. Although we do not have crystallographic evidence or single-particle cryo-electron microscopy data to confirm the molecular structure of CFAP61, our interactome data allow for an estimate of the position of CFAP61 in RS and CSC.

Dynein arms, DRC and RS regulate sperm motility(Viswanadha et al., 2017).Interestingly, deficiency of dynein arms, DRC or RS affect the length of the flagella and the assembly of microtubules and appendages in different degrees in spermatozoa(Ferheen Abbasi et al., 2018; Ben Khelifa et al., 2014; Dutcher et al., 2020; C. Liu et al., 2020b). The reasons behind this phenotype are not clear, but we observed that these abnormalities may be independent of motility regulation. For example, as a DRC component, *Tcte1* deletion affects sperm motility, but has little effect on sperm tail morphology(Castaneda et al., 2017). Here, axonemes from *Cfap61^−/−^* early round spermatids, appeared normal, however their structure became disordered in elongating spermatids and spermatozoa, but the positioning of the centrioles and of the implantation fossa formation were not affected. This abnormal phenomenon of spindle wire assembly cannot be explained only by the known functions of RS or CSC. In the early spermatids stage, the 12S RS complex contributes to form the axoneme, but is eliminated in *Cfap61^−/−^* elongating spermatid. These results suggest that CFAP61, is an important protein in the late stages of RS assembly and is necessary for the structural stability of the flagellum.

We observed that other proteins that interact with CFAP61 may also be involved in the assembly process of the flagellum and that IFT proteins were retained in the flagellum in *Cfap61^−/−^* elongating spermatids. The reason for IFT retention is still not clear, but may be due to the disorganization of the axoneme or could mean that the absence of CFAP61 has a direct effect on the IFT movement. However, no abnormal axoneme assembly was observed in the cilia of respiratory tract cells, suggesting that CFAP61 is not essential in these cells or that a stricter checkpoint mechanism may exist in the process of spermatozoa flagellum formation. In addition, we found that CFAP61 interacts with a large number of phosphorylation related proteins, although the specific functions and subcellular localization of these proteins are still unclear and need to be further studied.

In conclusion, we described significant differences in the assembly and structural stability of the axoneme in *Cfap61^−/−^* flagellum and cilia. Therefore, Cfap61 has a unique function in the process of flagellum formation. These results provide a theoretical basis for differential regulation of cilia/flagellum formation and MMAF physiopathology.

## Materials and Methods

### Study Patients

Here, we re-analyzed our data obtained by WES performed for a total of 167 individuals affected by primary infertility associated with a MMAF and described previously(Coutton et al., 2019). We focused the analyses on patients for whom no genetic diagnosis had been reached. All the recruited subjects displayed isolated infertility with no other clinical features; in particular, primary ciliary dyskinesia (PCD) syndrome was excluded. All individuals presented with a typical MMAF phenotype, which is characterized by severe asthenozoospermia (total sperm motility below 10%; normal values 40%) with at least three of the following flagellar abnormalities present in >5% of the spermatozoa: short, absent, coiled, bent, or irregular flagella. All individuals had a normal somatic karyotype (46, XY) with normal bilateral testicular size, hormone levels, and secondary sexual characteristics. Informed consent was obtained from all the individuals participating in the study according to local protocols and the principles of the Declaration of Helsinki. The study was approved by local ethics committees, and samples were then stored in the CRB Germethèque (certification under ISO-9001 and NF-S 96-900) according to a standardized procedure or were part of the Fertithèque collection declared to the French Ministry of health (DC-2015-2580) and the French Data Protection Authority (DR-2016-392).

### Whole exome sequencing (WES)

Genomic DNA was isolated from EDTA blood using the DNeasy Blood & Tissue Kits from QIAGEN SA (Courtaboeuf, France) or from saliva using the Oragen DNA extraction kit (DNAgenotech®, Ottawa, Canada) or from saliva using the Oragen DNA extraction kit (DNAgenotech®, Ottawa, Canada). Coding regions and intron/exon boundaries were sequenced on the Novogen platform (agilent v6, HiSeqX,) after enrichment with Agilent kits (Agilent Technologies, Wokingham, UK). Exomes data were analysed using a bioinformatics pipeline developed in-house using two modules, both distributed under the GNU General Public License v3.0 and available on github: https://github.com/ntm/grexome-TIMC-Primary and https://github.com/ntm/grexome-TIMC-Secondary and as described in part in reference(Arafah et al., 2020). Variants with a minor allele frequency greater than 1% in gnomAD v2.0, 3% in 1,000 Genomes Project phase 3, or 5% in NHLBI ESP6500, were filtered out and only variants predicted to have high-impact (e.g. stop-gain or frameshift variants) by variant Effect Predictor v92 (McLaren et al., 2016) were scrutinized.

### Animals

All mice used in this study were housed in a controlled environment at 20–22°C with a 12h light/dark cycle, 50–70% humidity, food and water *ad libitum*. All studies were approved by the Institutional Animal Care and Use Committees of Nanjing Medical University (Approval No. IACUC-1810020), Nanjing, China. All experiments with mice were conducted ethically according to the Guide for the Care and Use of Laboratory Animals and institutional guidelines.

### Quantitative RT-PCR assay

Total RNA was extracted from mouse tissues using Trizol reagent (ThermoFisher). cDNA synthesis was carried out on 1 μg of total RNA, using a HiScriptIII RT SuperMix (Vazyme, R323, Nanjing,China) according to the manufacturer’s instructions. The cDNA (dilution 1:4) was then analyzed by quantitative RT-PCR in a typical reaction of 20 μl containing 250 nmol/l of forward and reverse primers, 1 μl of cDNA and AceQ qPCR SYBR Green Master Mix (Vazyme, Q131, Nanjing, China). The reaction was initiated by preheating at 50°C for 2 min, followed by 95°C for 5 min and 40 amplification cycles of 10s denaturation at 95°C and 30s annealing and extension at 60°C. Gene expression was normalized to 18s rRNA within the log phase of the amplification curve. The primer sequences are listed in Supplementary file 1.

### Minigene splicing assay

Minigene splicing assay was carried out as described previously(Windpassinger et al., 2017).C20orf26 (NM_015585.4) minigenes containing exon2,a 357 bp fragment of the 5’ flanking intron, a 238 bp fragment of the 3’ flanking intron and differential for c.143+5 G>A mutation were amplified by PCR with oligos carrying the recombinant sites XhoI and BamHI. The PCR fragment was then cloned into the pET01 vector (MoBiTec, Goettingen, Germany). All minigene plasmids were sequenced to verify the correct insertion of mutated and wild-type DNA fragments. The splicing assay was performed by transiently transfecting HEK293T cells with each plasmid using Lipofectamine 2000 (ThermoFisher). At 48 hours post-transfection, cells were harvested and total RNA was extracted and reserve-transcribed. The resulting cDNAs were amplified by PCR with the forward primer corresponding to upstream exon A and the reverse primer complementary to downstream exon B. The primer sequences were as follows: Forward 5’-CCAGTTGAGGAGGAGAAC -3’and reserve 5’-CCAAGGTCTGAAGGTCAC -3’. PCR products were separated by 2% agarose gel electrophoresis and fragments were analyzed by Sanger sequencing.

### Antibodies

Rabbit antibodies specific for RSPH9 (23253-1-AP), NME5 (12923-1-AP), DNAJB13 (25118-1-AP), DYNLL2 (16811-1-AP), SLC2A3 (20403-1-AP) and IFT74 (27334-1-AP) were purchased from Proteintech (Rosemont, IL, USA). Rabbit antibody specific for β-Actin (ab8229) was purchased from Abcam. Mouse anti-FLAG M2 (F3165) and mouse anti-Acetylated Tubulin (T6793) were purchased from Sigma-Aldrich. The rabbit antibody specific for the DDDDK-tag (PM020) that was used for co-immunoprecipitation and the mouse antibody specific for HA-tag (M180-3) were purchased from Medical & Biological Laboratories (Nagoya, JP). Mouse anti-AKAP4 was purchased from BD Biosciences (California, USA).

The specific antibody for CFAP61 was generated according to the published method (M. Liu et al., 2014). Briefly, mouse CFAP61 (aa 223–348 and aa 1103-1230) was expressed as His fusion proteins in E.coli using the pET-28a(+) vector, then the fusion proteins were affinity purified with Ni-NTA His Bind Resin. Two rabbits were immunized with the fusion protein, respectively. The resulting working antisera are anti-CFAP61.

### Generation of *Cfap61^−/−^* Mice by CRISPR/Cas9

The *Cfap61* knockout mice were generated using CRISPR/Cas9 genome editing as described below. In brief, we selected two sgRNA targets to generate a deletion of the exon4 of *Cfap61* in mouse. The target sequences of sgRNA were 5’-GCTGTTTATGCACTTCTTTGTGG-3’ and 5’-GATTTCTTTGAGGCAGCCAGTGG -3’. The two complementary DNA oligos of each sgRNA target were annealed and ligated to the BsaI-digested pUC57-T7-sgRNA vector. The sgRNA templates were obtained from sgRNA plasmids by PCR amplification with primers Trans PCR For (5’-GAAATTAATACGACTCACTATAGG-3’) and Trans PCR Rev (5’-AAAAGCACCGACTCGGTGCCA-3’). Then, the PCR products were purified using a MinElute PCR Purification Kit (28004, QIAGEN). Two sgRNA were produced using the MEGAshortscript Kit (AM1354, Ambion) and purified using the MEGAclear Kit (AM1908, Ambion) according to the manufacturer’s instructions. The Cas9 plasmid (Addgene No. 44758) was linearized with AgeI and then purified using a MinElute PCR Purification Kit (28004, QIAGEN). Cas9 mRNA was produced by *in vitro* transcription using a mMESSAGE mMACHINE T7 Ultra Kit (AM1345, Ambion) and purified using a RNeasy Mini Kit (74104, QIAGEN) per the manufacturer’s instructions. Mouse zygotes were coinjected with Cas9 mRNA (50 ng/μL), sgRNA (20 ng/μL). The injected zygotes were transferred into pseudo-pregnant recipients. The newborn mice (7 days old) were tagged by a toe cut, and DNA was extracted using the Mouse Direct PCR Kit (B40013, Biotool). PCR amplification was carried out with primers (Forward: 5’-AGGCAGTGAGTGAAGTGT -3’, Reverse: 5’-TAAGTTGGCGAGGCTTGA -3’) using PrimeSTAR HS DNA Polymerase (DR010A, Takara) under the following conditions: 95°C for 5 min; 35 cycles of 95°C for 30 s, 62°C (–0.2°C/cycle) for 30 s, and 72°C for 30 s; and a final step of 72°C for 5 min. The PCR products were subjected to Sanger sequencing.

### Fertility Test

Adult mice from each genotype were subjected to fertility tests in which each male was mated with three wild-type C57BL/6 female mice, and the vaginal plug was checked every morning. The dates of birth and number of pups in each litter were recorded.

### Sperm Analysis

Epididymal sperm was obtained by making small incisions throughout the cauda of the epididymis, followed by extrusion and suspension in human tubal fluid culture medium (In Vitro Care, Frederick, MD, USA) supplemented with 10% FBS at 37 °C. Sperm samples (10 μl) were used for computer-assisted semen analysis (Hamilton-Thorne Research, Inc., Beverly,MA, USA). The remaining of sperm samples were fixed in 4% paraformaldehyde for 30min and subsequently spread on slides. The H&E staining was conducted using standard methods for sperm morphology study. Over 200 spermatozoa were examined, and morphological abnormalities were evaluated as described previously (Li, Wu, Li, Tian, Kherraf, Zhang, Ni, Lv, Liu, Tan, Shen, Amiri-Yekta, Cazin, Zhang, et al., 2020) following the WHO guidelines. Each spermatozoon was classified in only one morphological category according to its major flagellar abnormality.

### Histological Analysis

Mouse testes, epididymis and tracheas were collected from at least three mice for each genotype. The testes and epididymis were fixed in modified Davidson’s fluid for up to 24 h while tracheas were fixed in 4% paraformaldehyde in PBS overnight and stored in 70% ethanol. The samples were then dehydrated through a graded ethanol series and embedded in paraffin. Tissue sections (thickness 5μm) were prepared and mounted on glass slides and HE staining was performed according to standard procedures. Periodic Acid-Schiff (PAS) staining was carried out using the Sigma Aldrich PAS staining kit (395B).

### Transmission electron microscopy (TEM)

Ultrastructural examination were performed as described below. Briefly, the epididymal sperm was fixed with 2.5% glutaraldehyde overnight,postfixed with 2% OsO4 and embedded in Araldite. Ultrathin sections (80 nm) were stained with uranyl acetate and lead citrate and analyzed by an electron microscope (JEM.1010, JEOL).

### Scanning electron microscopy (SEM)

Spermatozoa and trachea from normal and *Cfap61-*mutated male mice were fixed in 2.5% phosphate-buffered glutaraldehyde at 4°C for 2 hours. Immobilized spermatozoa were deposited on poly-L-lysine coated coverslips. Subsequently, spermatozoa and trachea were washed in PBS solution, dehydrated via an ascending gradient of cold 30%, 50%, 70%, 80%, 90% and 100% ethanol, and dried at critical point using a Leica EM CPD300 Critical Point Dryer (Wetzlar, Germany). Specimens were then attached to specimen holders and coated with gold particles using an ion sputter coater (EM ACE200, Leica, Wetzlar, Germany) before being viewed with a Helios G4 CX scanning electron microscope (Thermo scientific).

### Fractionation of spermatozoa

Sperm protein fractionation was performed as described previously (Castaneda et al., 2017). Spermatozoa were suspended in 1% TritonX-100 lysis buffer (50 mM NaCl, 20 mM Tris·HCl, pH 7.5, protease inhibitor mixture) and incubated at 4 °C for 2 h. The sample was centrifuged at 15,000 × g for 10 min to separate the Triton-soluble fraction (supernatant) and the Triton-resistant fraction (pellet). The pellet was resuspended in 1% SDS lysis buffer (75 mM NaCl, 24 mM EDTA, pH 6.0) and incubated at room temperature for 1 h. The sample was centrifuged at 15,000 × g for 10 min to separate SDS-soluble fraction (supernatant) and SDS-resistant fraction (pellet). The pellet was dissolved in sample buffer and boiled for 10 min.

### Western blot analysis

Western blotting was performed as described below. Briefly, protein extracts were prepared using lysis buffer (8 M urea, 50mM Tris-HCl pH 8.2, 75Mm NaCl) in the presence of 1x cOmpleteTM EDTA-free Protease Inhibitor Cocktail (Roche, Basel, Switzerland). The proteins were separated by SDS-PAGE and transferred onto a polyvinylidene difluoride membrane. The membrane was blocked with 5% non-fat milk in TBS solution for 2 h at room temperature and incubated overnight at 4°C with primary antibodies. The membranes were washed with TBST (0.1% Tween-20 in TBS) buffer three times and incubated at room temperature for 2 h with secondary antibodies. The signals from the detected proteins were visualized using SuperSignal West Femto Chemiluminescent Substrate (Thermofisher).

### Plasmids construction

Full-length cDNA encoding Cfap61, Rsph3a, Rsph9, Ropn1, Ropn1l, Caml4, Armc4, Dynll2, Maats1, Dynltla, Dyncli2, Dynll1, Dynlrb2, Ift22, Ift74, Ift81 and Wdr35 were amplified by PCR with oligos carrying the recombinant sites and cloned into pcDNA3.1(+) vector (Thermofisher) in which a FLAG or HA epiptope was introduced prior to the multicloning site. The primers used to amplify each gene are listed in Supplementary file 2. Chlamydomonas genes were chemically synthesized by GenScript (Nanjing,China) and inserted individually into vector plasmid pcDNA3.1(+)-N-HA or pcDNA3.1(+)-N-FLAG.

### Cell culture

HEK293T cells were maintained in DMEM high glucose supplemented with 10% fetal bovine serum (Gibco), Penicillin-Streptomycin (100U/ml,Thermofisher). Transfections of HEK293T cells were performed using using Lipofectamine 2000 (11668019, ThermoFisher) according to manufacturer’s instructions.

### Immunoprecipitation

Two days after transfection, cells were lysed with RIPA Lysis Buffer (P0013C, Beyotime) supplemented with 1x cOmpleteTM EDTA-free Protease Inhibitor Cocktail (Roche, Basel, Switzerland) for 40 min at 4 °C and then clarified by centrifugation at 12,000 × g for 20 min. The lysates were precleared with 10µL ProteinA magnetic beads (10008D, ThermoFisher) for 1 h at 4 °C. Precleared lysates were incubated overnight with Anti-DDDDK-tag at 4 °C. Lysates were then incubated with 50 μL Protein A magnetic beads for 4h at 4 °C. The beads were washed three times with RIPA Lysis Buffer and boiled for 5min in SDS loading buffer before SDS/PAGE.

CFAP61 was immunoprecipitated from mouse testis using the Pierce crosslink IP kit (Thermo Scientific) with anti-CFAP61 antibody described above, testis from knock-out animals was used as negative control. IP was performed according to the manufacturer’s instructions.

### Mass spectrometry

Eluates were precipitated with five volumes of −20°C pre-chilled acetone followed by trypsin digestion. LC-MS/MS analysis was performed on EASY-nanoLC 1000 system (Thermo Scientific) coupled to an Orbitrap Fusion Tribrid mass spectrometer (Thermo Scientific) by a nano spray ion source. Tryptic peptide mixtures were injected automatically and loaded at a flow rate of 20 μl/min in 0.1% formic acid in LC-grade water onto an analytical column (Acclaim PepMap C18, 75 μm x 25 cm; Thermo Scientific). The peptide mixture was separated by a linear gradient from 5% to 38% of buffer B (0.1% formic acid in ACN) at a flow rate of 300 nl/min over 53 minutes. Remaining peptides were eluted by a short gradient from 38% to 90% buffer B in 1 minutes. Analysis of the eluted peptides was done on an Orbitrap Fusion Tribrid mass spectrometer. From the high-resolution MS pre-scan with a mass range of 335 to 1400, the most intense peptide ions were selected for fragment analysis in the orbitrap depending by using a high speed method if they were at least doubly charged. The normalized collision energy for HCD was set to a value of 28 and the resulting fragments were detected with a resolution of 120,000. The lock mass option was activated; the background signal with a mass of 445.12003 was used as lock mass. Every ion selected for fragmentation was excluded for 30 seconds by dynamic exclusion. Data were processed with MaxQuant software (version 1.6.10.43) and Mouse reference proteome from SwissProt database (release 2019_07) using standard parameters. The mass spectrometry proteomics data have been deposited to the ProteomeXchange Consortium via the PRIDE(Perez-Riverol et al., 2019) partner repository with the dataset identifier PXD024469.

### Immunofluorescence

For testis and trachea cryosections, immunostaining was performed as previously described(Castañeda et al., 2014). For spermatozoa, samples were obtained as described above. For germ cells, samples were squeezed out from the seminiferous tubules onto slide glasses and air-dried at room temperature. The samples were then fixed with 4% paraformaldehyde in PBS for 30 min. After three 10-min washes with PBS, heat-induced antigen retrieval was carried out by boiling the slides in 10 mM citrate buffer (pH 6.0) in a microwave oven for 10 min. After three 10-min washes with PBST (0.1% Triton X-100 in PBS), the slides were blocked with 5% BSA diluted in PBST for 1 h and then incubated with primary antibodies at 4°C overnight. After incubation with the secondary antibody at room temperature for 2 h, the slides were incubated with Hoechst 33342 for 5 min. Finally, the slides were washed in PBS and then mounted with VectaShield or Immu-Mount. Slides were viewed with an LSM800 confocal microscope (Carl Zeiss AG) or TCS SP8X confocal microscope (Leica Microsystems).

### Recording of cilia motility in trachea

Mouse trachea were removed by dissection and placed in DMEM high glucose supplemented with 10% fetal bovine serum (Gibco). Trachea were opened on the dorsal side and cut into 5 mm squares under a stereoscopic microscope. The tissue pieces were observed in a confocal dish (BDD012035, BIOFIL) on a glass slide (801011, NEST) with a scotch tape spacer under a 40× objective (CFI S Plan Flour ELWD NAMC) of an inverted microscope (Eclipse Ti2-U, Nikon).

### Statistical analysis

All experiments were repeated at least three times. The differences between treatment and control groups were analyzed using one-way ANOVA or unpaired two-tailed t-tests. P-values < 0.05 were considered statistical significance. All data represent the mean ± the standard error of the mean. Analyses were performed using the Microsoft Excel or GraphPad Prism 6.0.

## Acknowledgements

This work was supported by the National Key Research and Development Program of China 2016YFA0500902 (to M.L.); Natural Science Foundation of China (32070842, 31771654 and 31530047 to M.L.); the Natural Science Foundation of Jiangsu Province (Grants No. BK20190081 to M.L.); and Qing Lan Project (to M.L.); the FLAGEL-OME project (to P. F. Ray) : ANR-19-CE17-001.

## Author contributions

S.L., P.F.R., and M.L. initiated the project and designed the experiments; S.L., P.F.R., and M.L.wrote the paper. S.L. performed most of the experiments and analysis; J.Z., Z.E.K., S.S., X.Z., C.Ca., C.Co., R.Z., S.Z., F.H., S.F.B.M., and C.A. performed some of the experiments and analysis; All authors read and approved the final manuscript.

## Ethics Declaration

Informed consent was obtained from all the individuals participating in the study according to local protocols and the principles of the Declaration of Helsinki. The study was approved by local ethics committees, and samples were then stored in the CRB Germethèque (certification under ISO-9001 and NF-S 96-900) according to a standardized procedure or were part of the Fertithèque collection declared to the French Ministry of health (DC-2015-2580) and the French Data Protection Authority (DR-2016-392).

All anmial studies were approved by the Institutional Animal Care and Use Committees of Nanjing Medical University (Approval No. IACUC-1810020), Nanjing, China. All experiments with mice were conducted ethically according to the Guide for the Care and Use of Laboratory Animals and institutional guideline.

---

**Figure 3-figure supplement 1:**
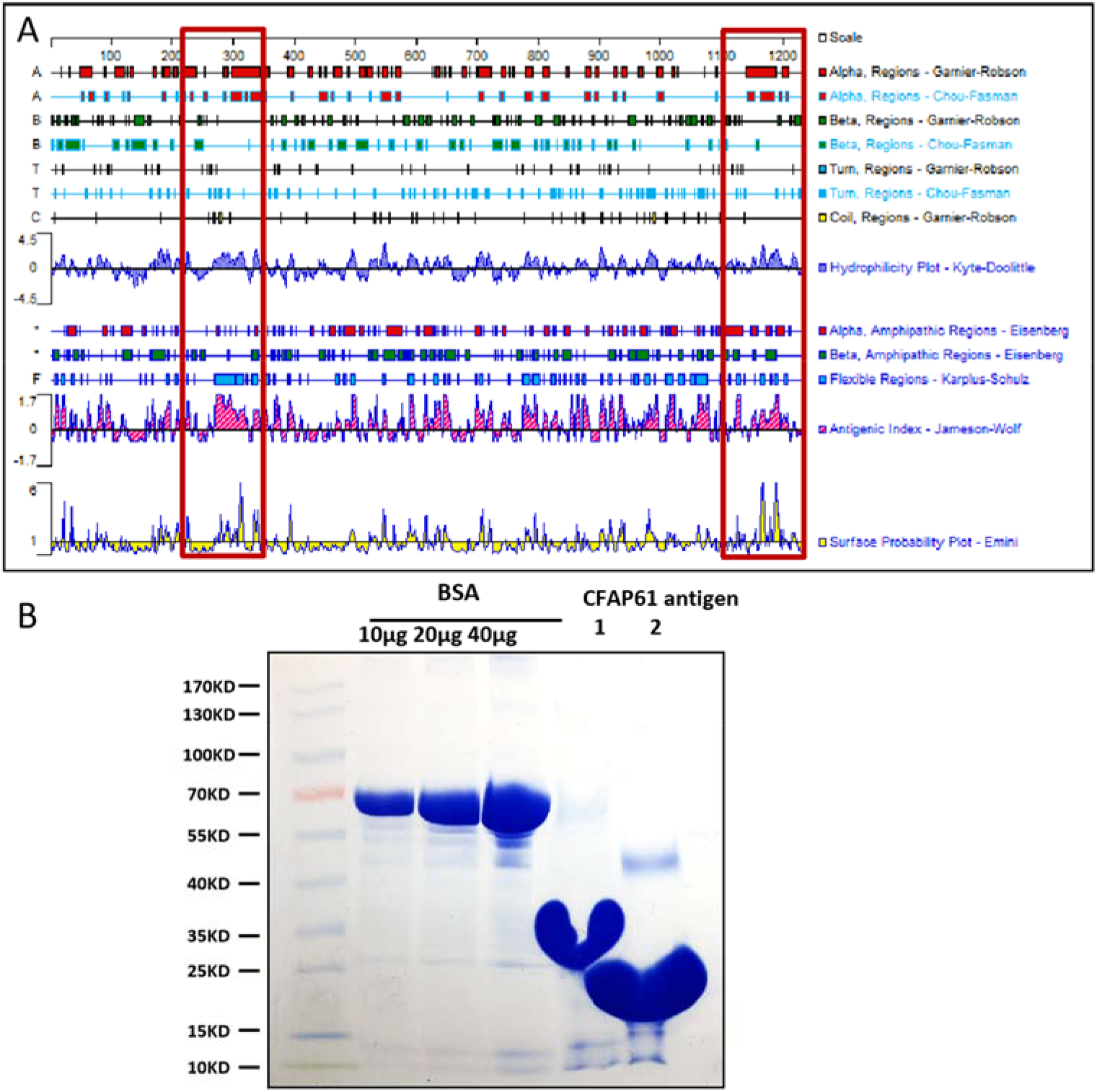
Antigen selection and purification. (A) Epitope prediction of CFAP61 by analyzing secondary structures and other indicators (flexibility, hydrophilicity, antigenicity, and surface probability). (B) Coomassie brilliant blue stained SDS-PAGE of selected antigens. Protein standards indicate molecular size in kilodaltons in leftmost lane, while next three lanes show the indicated microgram amounts of pure bovine serum albumin (BSA) and final two lanes show purified CFAP61 antigens.

**Figure 3-figure supplement 2:**
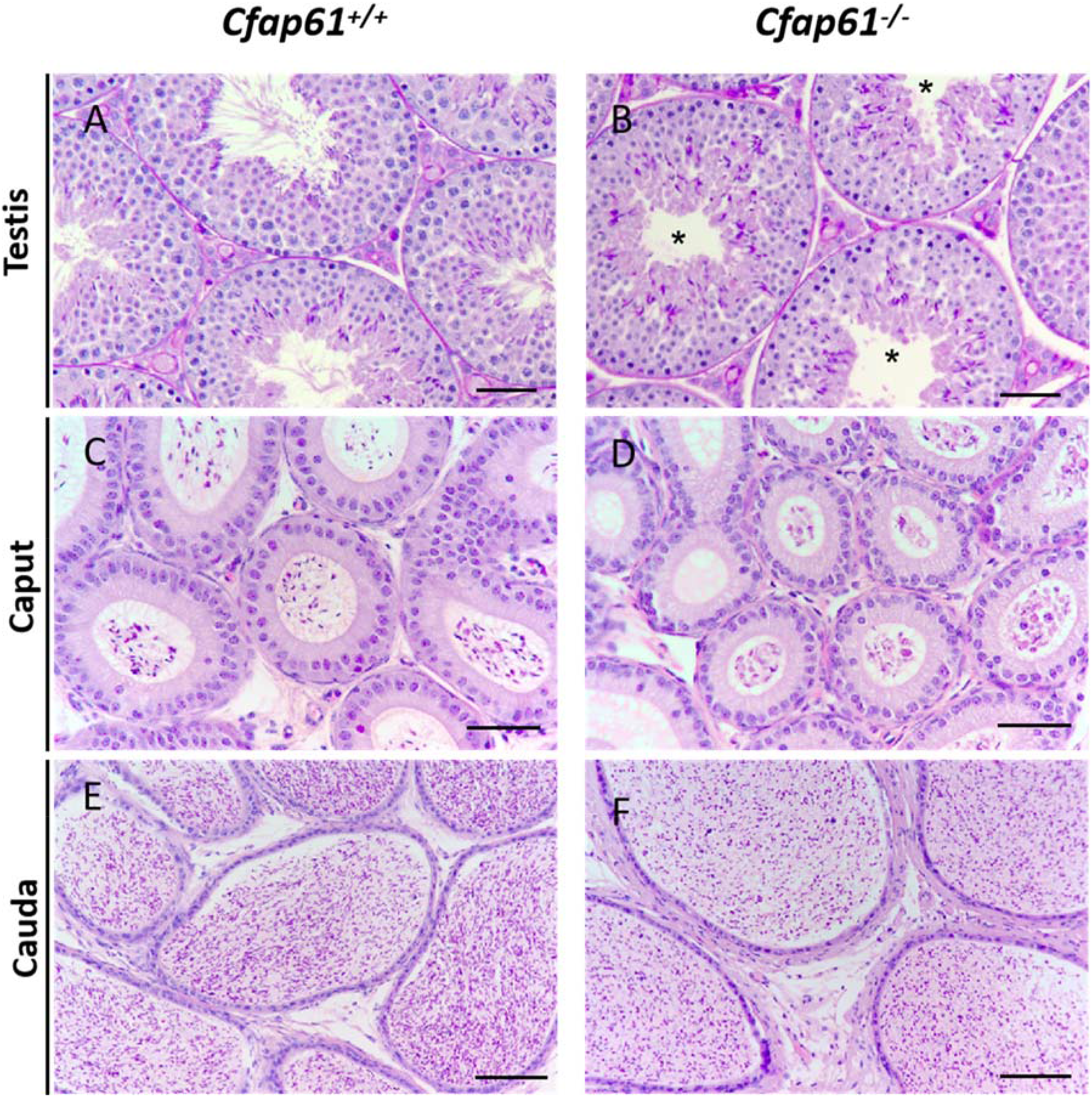
Spermatogenesis is abnormal in *Cfap61^−/−^* mice. Sections of periodic acid Schiff-stained testis and hematoxylin and eosin-stained caput and cauda epididymis from wild-type (A, C, E) and *Cfap61^−/−^* (B, D, F) mice.(A-D) Scale bars =50 μm.(E, F) Scale bars =100 μm.

**Figure 5-figure supplement 1:**
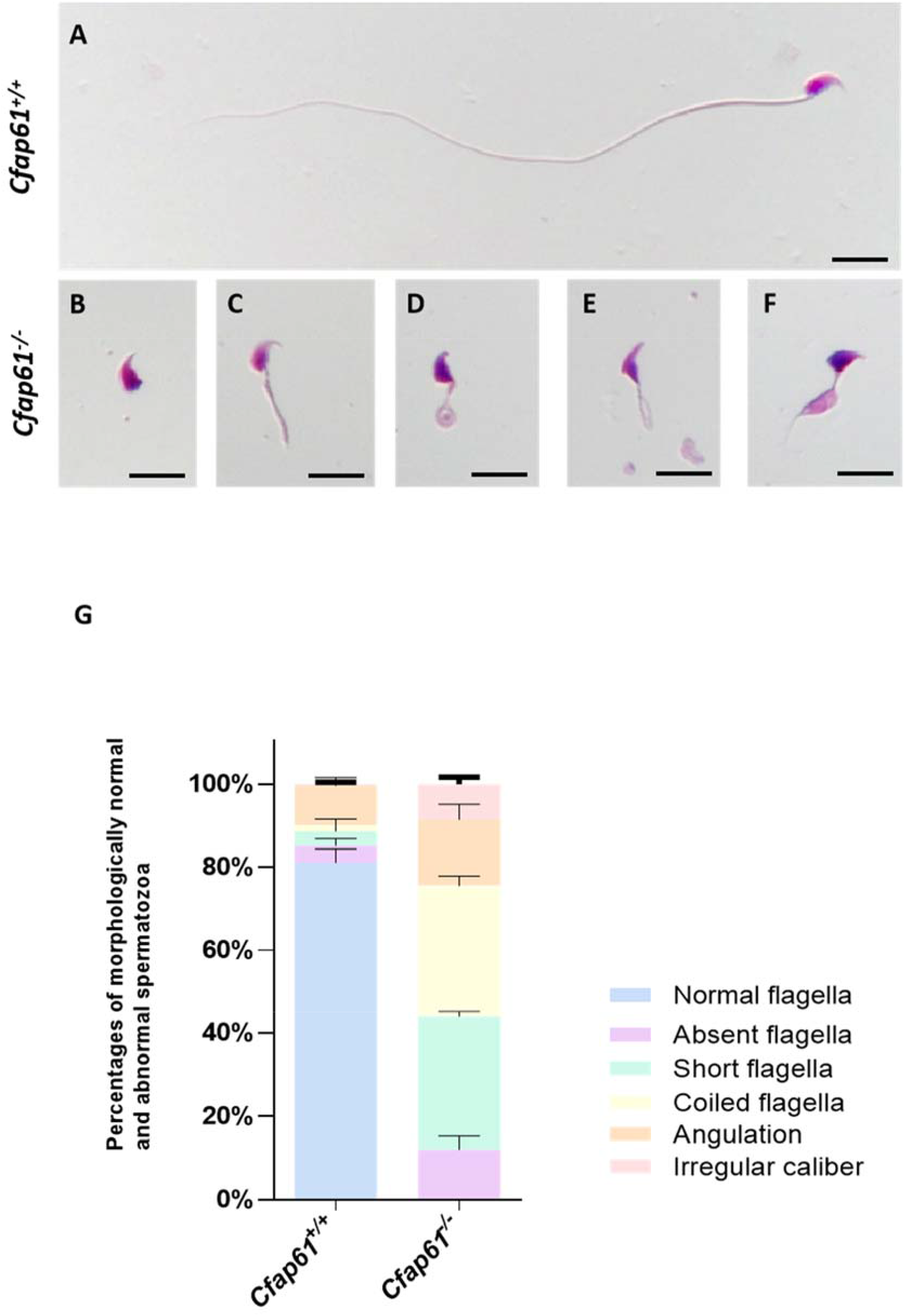
Sperm morphology study in *Cfap61^−/−^* male mice. (A) Light microscopy showed wild type spermatozoon with normal morphology while spermatozoa from *Cfap61^−/−^* mice manifested aberrant flagellar morphologies: absent (B), short (C), coiled (D), angulation (E) and irregular caliber (F), consistent with the MMAF phenotype. Scale bars =10 μm. (G) Percentage of morphologically normal and abnormal spermatozoa in WT and *Cfap61^−/−^* male mice (n=3), evaluated following the WHO guidelines. The left column shows that the percentage of spermatozoa with normal flagella(81.165± 1.895%), absent flagella(4.028±0.966%), short flagella (3.414±1.709%), coiled flagella (1.604±0.783%), anjulation (9.366±1.139%) and irregular caliber (0.423±0.259%) in the normal group.The right column shows that the percentage of spermatozoa with normal flagella(0%), absent flagella(12.085±1.852%), short flagella (31.969±0.654%), coiled flagella (31.542±1.317%), anjulation (15.833±2.125%) and irregular caliber (8.570± 0.940%) in the *Cfap61^−/−^* group.P_normal flagella_ (*Cfap61^−/−^* vs WT)= 1.77×10^^−6^, P_absent flagella_ (*Cfap61^−/−^* vs WT)= 0.0182, P_short flagella_ (*Cfap61^−/−^* vs WT)= 9.84×10^^−5^, P_coiled flagella_ (*Cfap61^−/−^* vs WT)= 4.04×10^^−5^, P_angulation_ (*Cfap61^−/−^* vs WT)= 0.055. P_irregular caliber_ (*Cfap61^−/−^* vs WT)= 0.001. Data represent mean±SEM.

**Figure 6-figure supplement 1:**
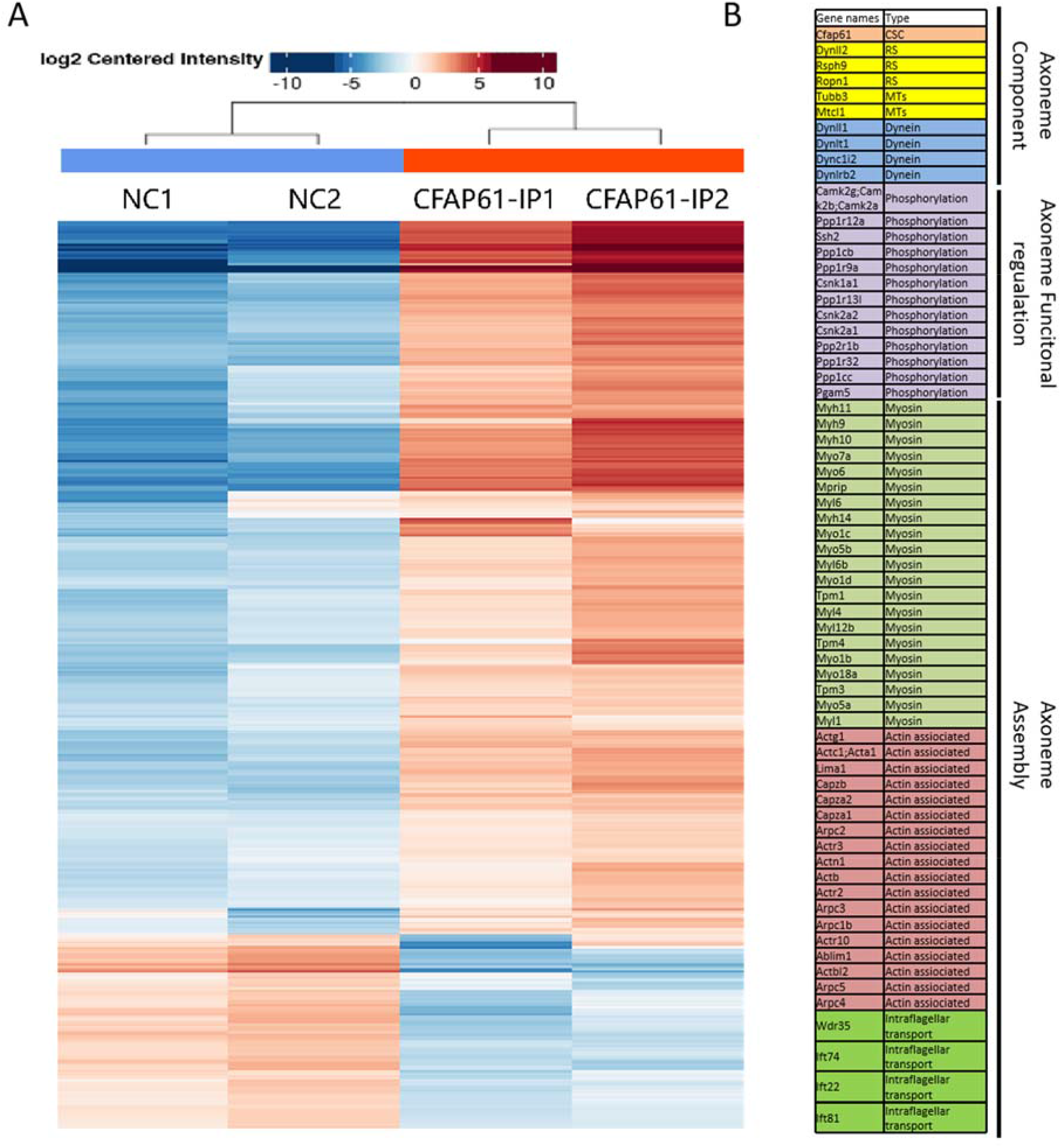
Identification of CFAP61 interacting proteins. (A) Heatmap representation of CFAP61 interacting proteins identified with anti-CFAP61, with pre-immune rabbit serum used as negative control. IP1-NC1 and IP2-NC2 represent data from two biological replicates. Data are available via ProteomeXchange with identifier PXD024469. (B) Functional classes of CFAP61 interacting proteins.

**Figure 6-figure supplement 2:**
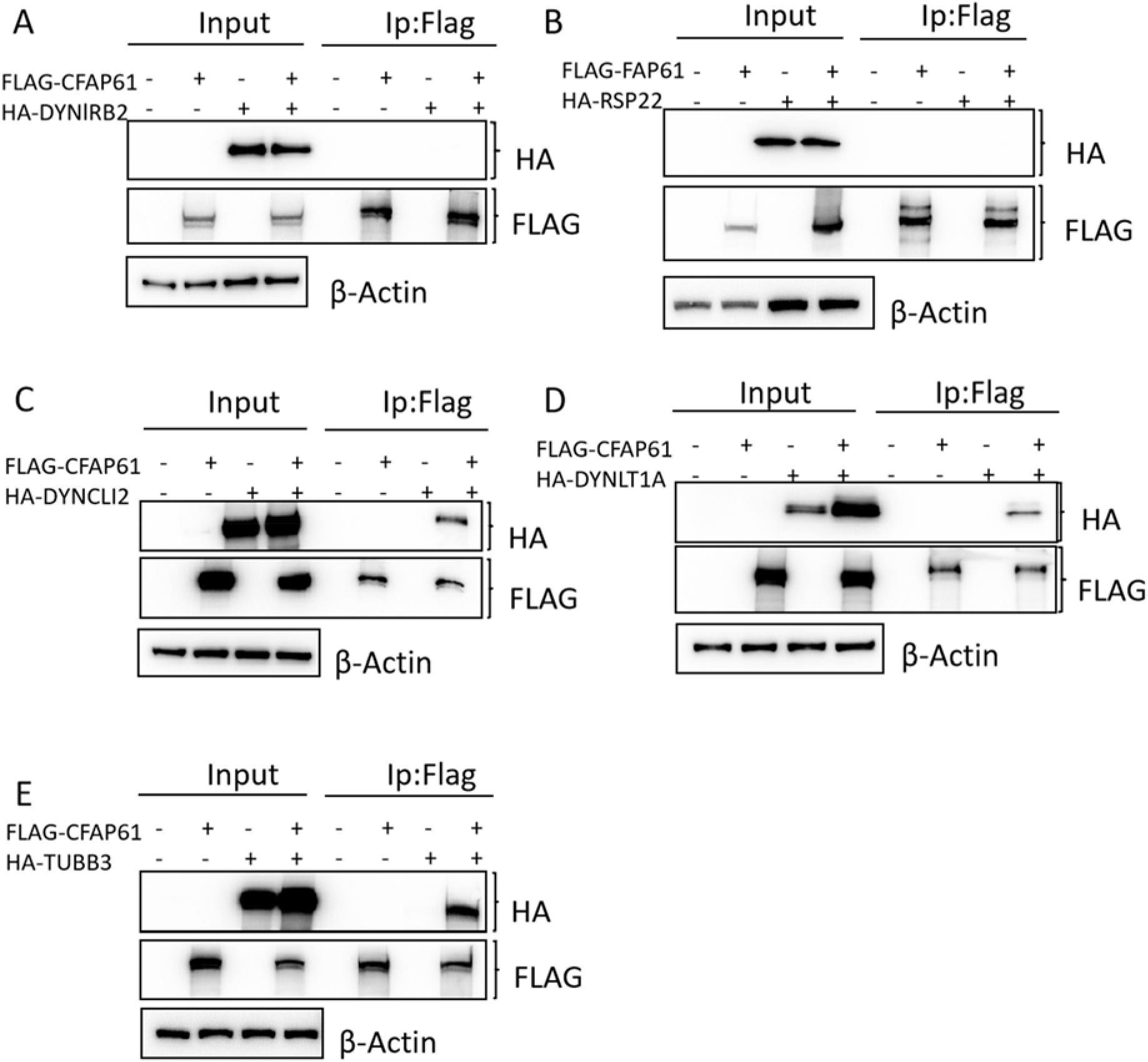
CFAP61 interacts with DYNEIN and TUBULIN. (A) Chlamydomonas RSP22 was expressed or co-expressed with FAP61 in HEK293T cells and FAP61 interaction with RSP22 was examined by co-immunoprecipitation.(B-E) Mouse dynein subunits (B-D) or TUBB3 (E) were expressed or co-expressed with CFAP61 in HEK293T cells and CFAP61 interaction with DYNLRB2,DYNCLI2,DYNLT1A,TUBB3 was examined by co-immunoprecipitation.

**Movie 1. Motility and movement of the sperm fr0om the caput epididymis of wild-type male mice.**

**Movie 2. Motility and movement of the sperm from the caput epididymis of *Cfap61^−/−^* male mice.**

**Movie 3. Ciliary motility of wild type tracheal epithelial cells. Movie 4. Ciliary motility of *Cfap61^−/−^* tracheal epithelial cells.**

**Supplementary file 1. Primer sequences For qRT-PCR**

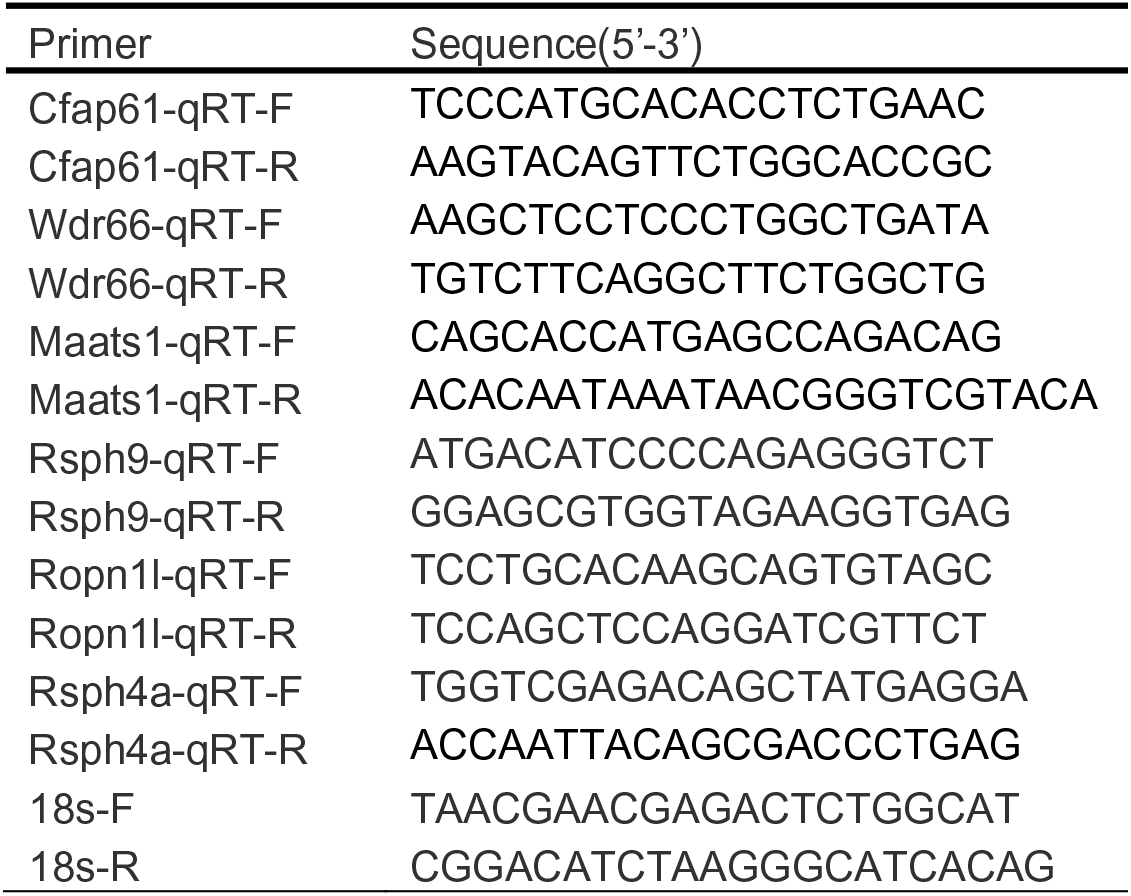

**Supplementary file 2. Primer sequences For plasmids construction**

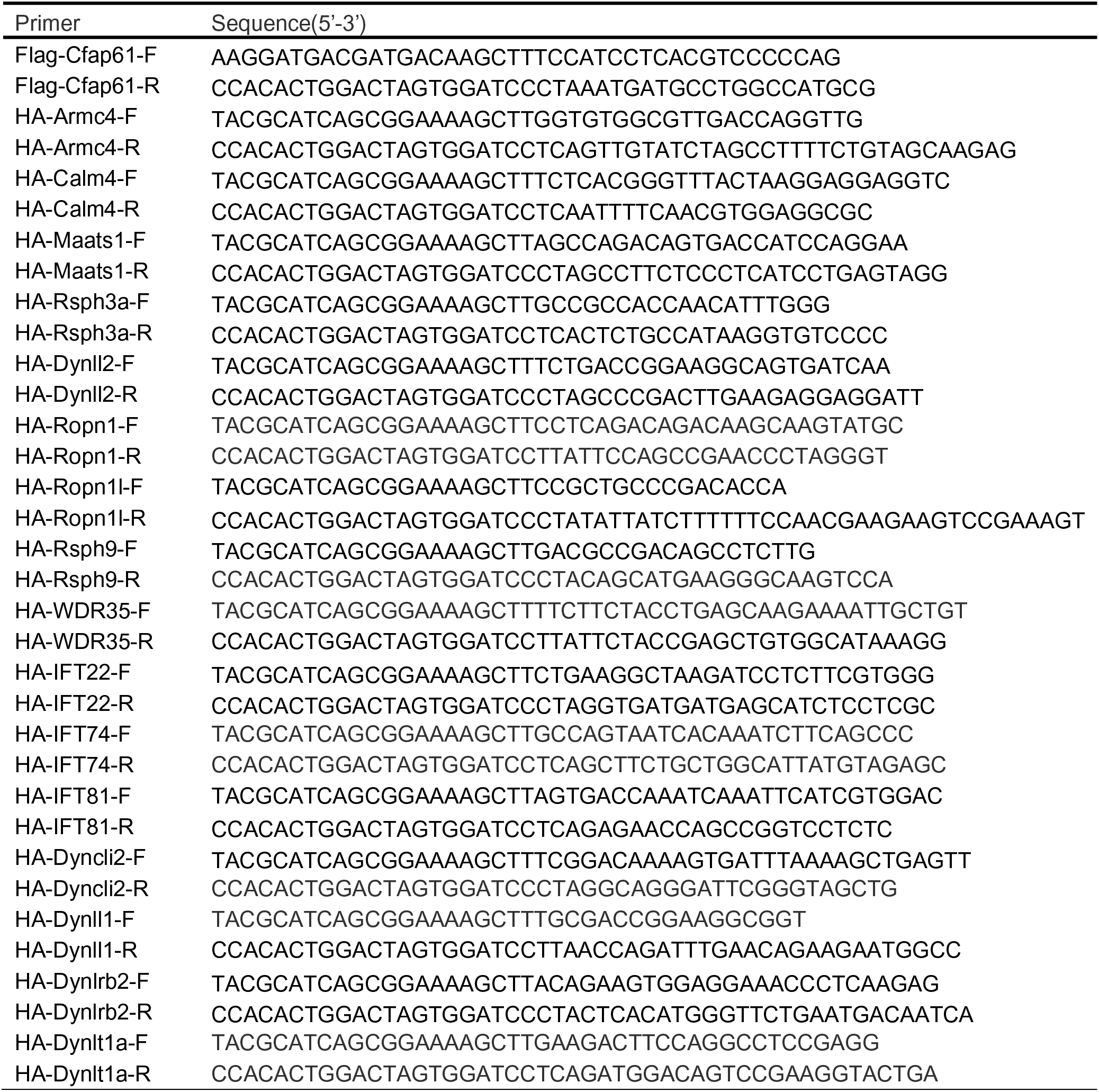

